## Supplementary Information for "Antibody-drug conjugates with dual payloads for combating breast tumor heterogeneity and drug resistance"

#### 1. SUPPLEMENTARY FIGURES AND TABLES

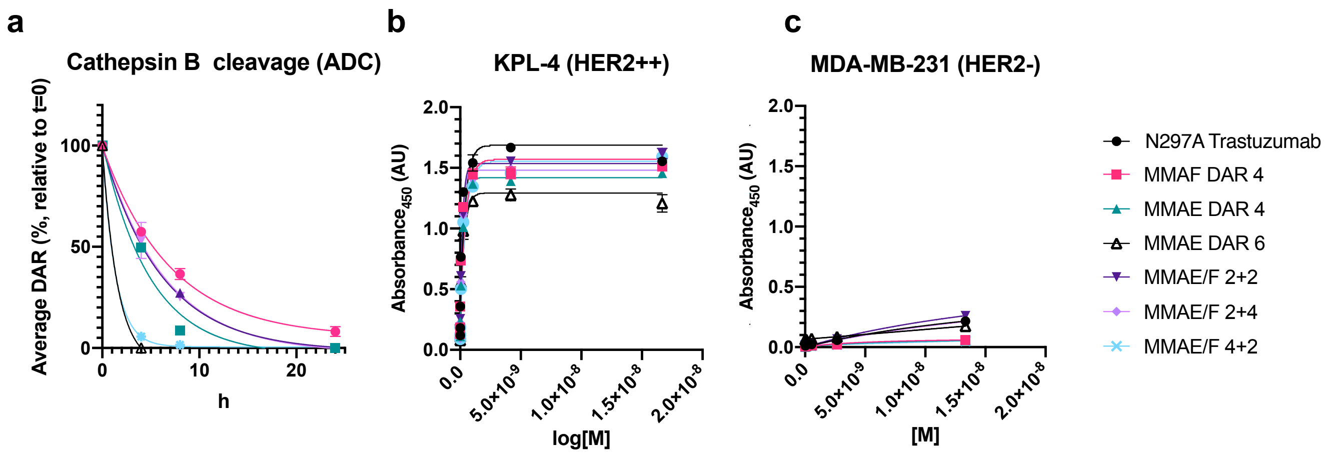

**Supplementary Fig. 1 Cathepsin B cleavage and HER2 binding assays. a**, Human cathepsin B-mediated cleavage of ADCs at 37 °C. The degree of loss of payload in each ADC was determined by HPLC. All assays were performed more than twice in technical duplicate. Error bars represent s.e.m. (*n* = 2). **b**, **c**, Saturation-binding curves obtained by cell-based ELISA. All assays were performed in triplicate and error bars represent s.e.m. The unmodified N297A anti-HER2 mAb and ADCs bound to KPL-4 cells (HER2 positive, **b**) with comparable binding affinities but not to MDA-MB-231 cells (HER2 negative, **c**).

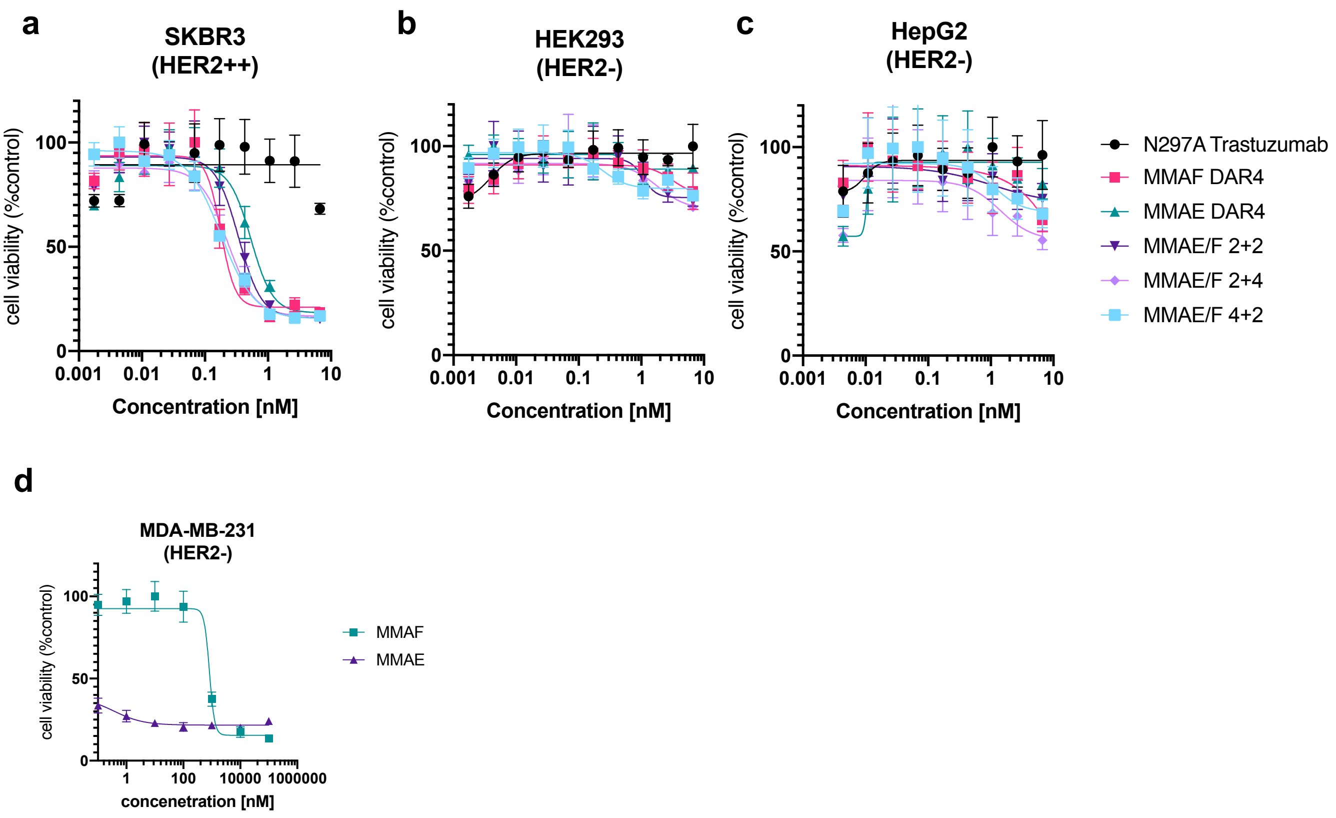

**Supplementary Fig. 2 In vitro cytotoxicity of ADCs**. **a**–**c**, Cytotoxicity of unconjugated N297A anti-HER2 mAb (black), MMAF DAR4 single-drug ADC (magenta square), MMAE DAR4 single-drug ADC (green triangle), MMAE/F 2+2 dual-drug ADC (dark purple inverted triangle), MMAE/F 2+4 dual-drug ADC (light purple diamond), and MMAE/F 4+2 dual-drug ADC (cyan square) in (**a**) SKBR-3, (**b**) HEK293, and (**c**) HepG2 cells. **d**, Cell killing potency of unconjugated MMAE (green square) and MMAF (purple triangle) in MDA-MB-231 cells. All assays were performed in quadruplicate. Error bars represent s.e.m.

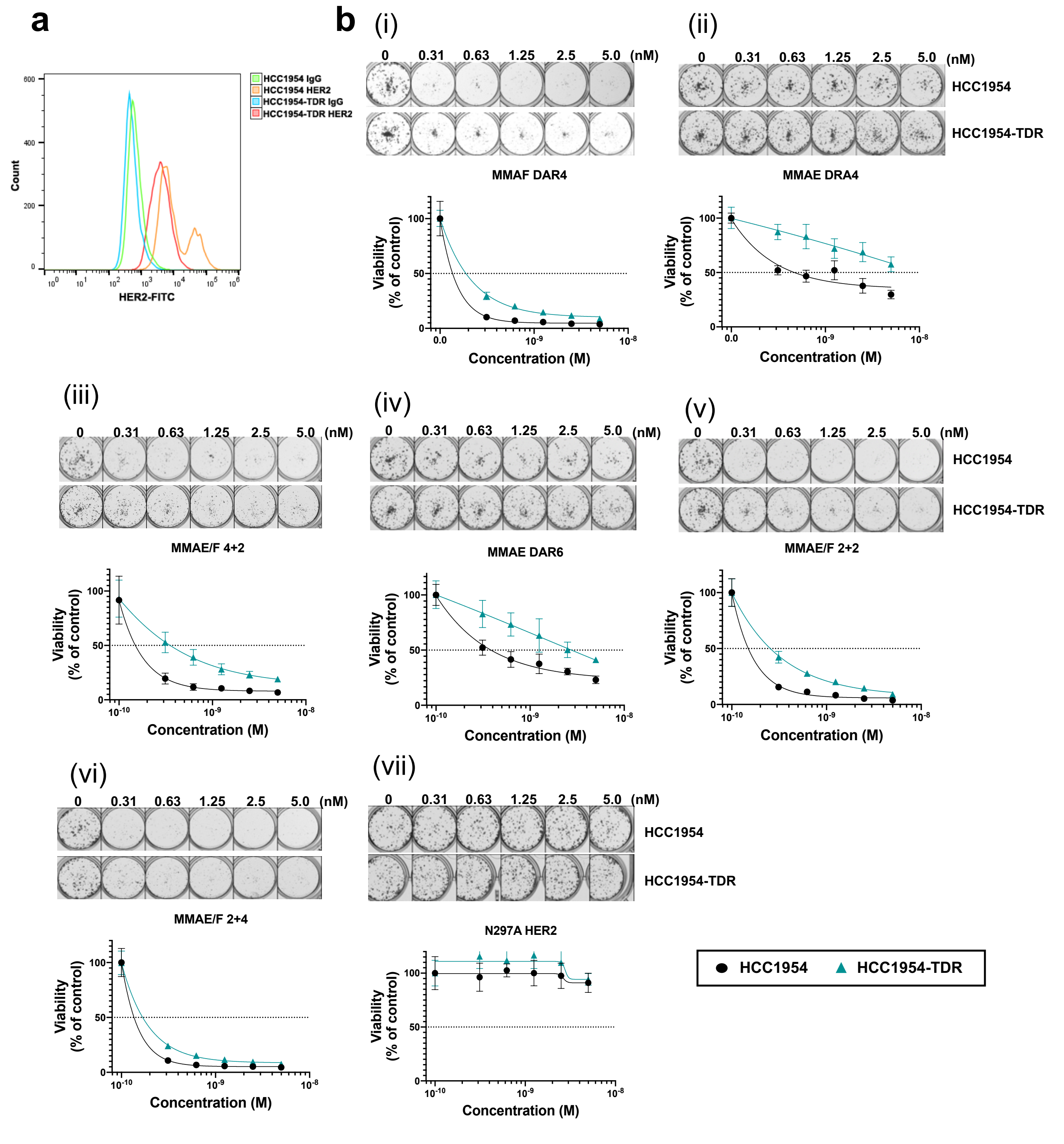

**Supplementary Fig. 3 HER2 expression and cell killing potency in HCC1954 and HCC1954-TDR cells. a**, flow cytometry analysis of HCC1954 and HCC1954-TDR cells labeled with anti-HER2 antibody or isotype control. HCC-1954-TDR shows a slightly reduced HER2 expression level whereas a HER2-high population exists in the parent HCC-1954 cells. **b**, Clonogenicity assay. We tested MMAF DAR4 single-drug ADC, MMAE DAR4 single-drug ADC, MMAE DAR6 single-drug ADC, MMAE/F 2+2 dual-drug ADC, MMAE/F 2+4 dual-drug ADC, MMAE/F 4+2 dual-drug ADC, and unconjugated N297A anti-HER2 mAb in HCC1954 (black) and HCC1954-TDR cells (green). All assays were performed in quadruplicate. Error bars represent s.d.

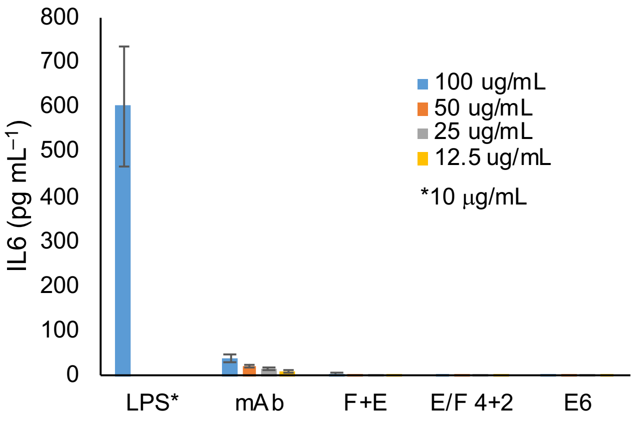

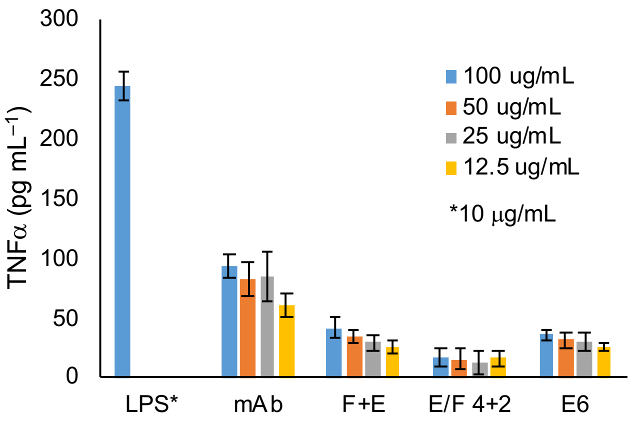

**Supplementary Fig 4 Cytokine release assay using primed THP-1 cells. a**,**b**, Levels of IL-6 (**a**) and TNF-α (**b**) released to cell culture supernatants were measured by ELISA after 24 h of incubation with N297A anti-HER2 mAb, 1:1 mixture of MMAF DAR 4 and MMAE DAR 4 single-drug ADCs (F+E), MMAE/F 4+2 dual-drug ADC (E/F 4+2), and MMAE DAR 6 single-drug ADC (E6) at 12.5–100 µg mL$^{-1}$. LPS (10 µg mL$^{-1}$) was used as a positive control. Bars represent mean ± s.d.

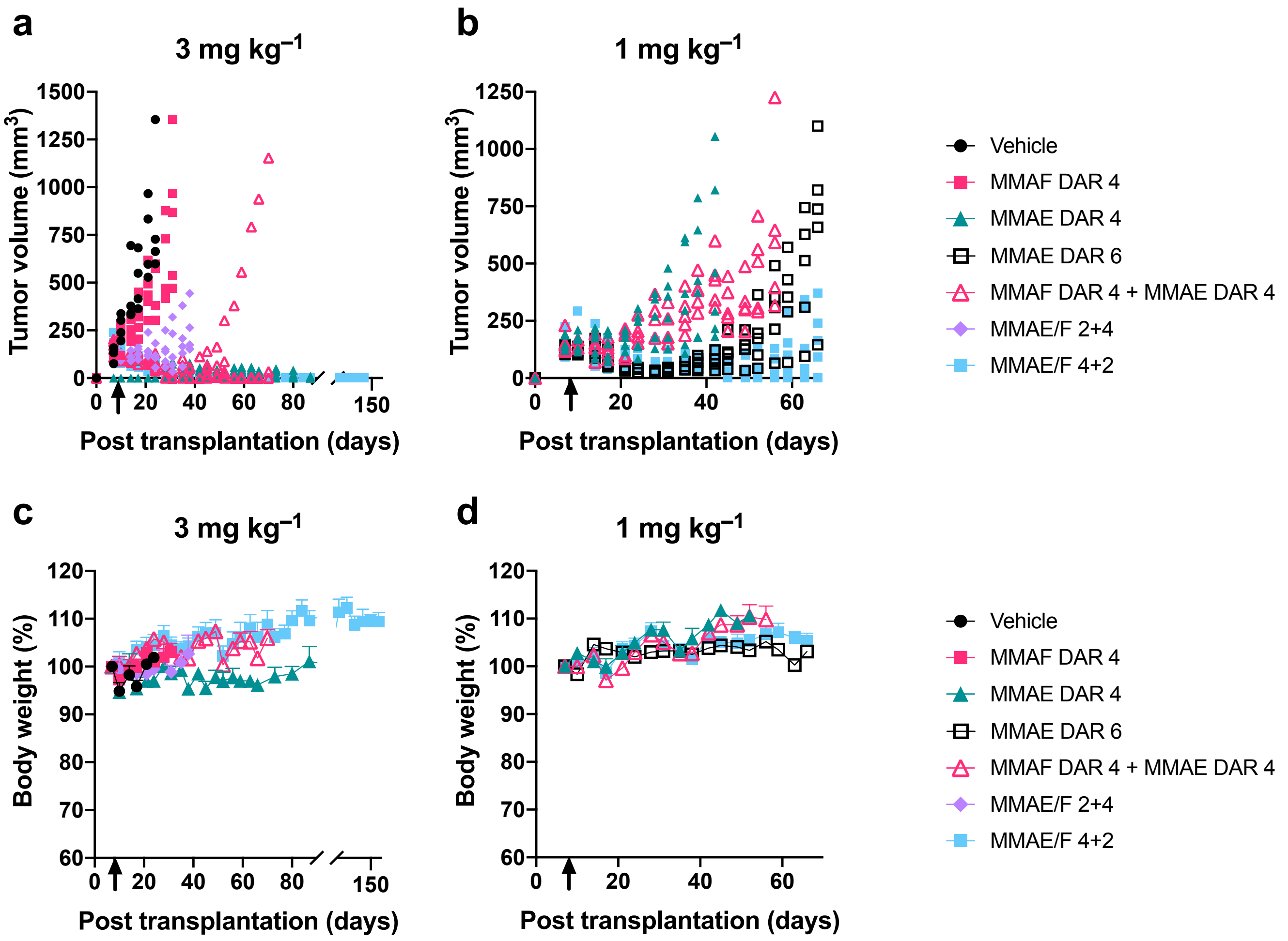

**Supplementary Fig 5 Antitumor activity in the JIMT-1/MDA-MB-231 (4:1) admixed xenograft tumor model and body weight change. a**,**b**, Tumor volume after treatment with each ADC at 3 mg $kg^{-1}$ (**a**) and 1 mg $kg^{-1}$ (**b**). Each dot represents an individual mouse. Figures generated using the same dataset are shown in the main article. Six–eight weeks old female NU/J nude mice with xenografted tumors were injected intravenously with a single dose of MMAF DAR 4 ADC (magenta square), MMAE DAR 4 ADC (green triangle), MMAE DAR 6 ADC (black blank square), a mixture of MMAF DAR 4 and MMAE DAR 4 ADCs (magenta blank triangle, 3 mg $kg^{-1}$ each or 1 mg $kg^{-1}$ each), MMAE/F 2+4 ADC (light purple circle), MMAE/F 4+2 ADC (cyan square) or vehicle control (black circle) at day 8 post transplantation (indicated with a black arrow). $n = 4$ for vehicle and MMAE DAR 4 ADC at 3 mg $kg^{-1}$; $n = 6$ for MMAE DAR 4 ADC at 1 mg $kg^{-1}$; $n = 5$ for all other groups. **c**,**d**, Body weight change after treatment with each ADC at 3 mg $kg^{-1}$ (**c**) and 1 mg $kg^{-1}$ (**d**). No significant body weight loss caused by either ADC was observed over the course of study. Error bars represent s.e.m.

**Supplementary Table 1 $K_D$ values in breast cancer cell lines (*n* = 3).** Values in parentheses are 95% confidential intervals.

| | $K_D$ (nM) | |
|---|---|---|
| | KPL-4 | MDA-MB-231 |
| Anti-HER2 mAb | 0.081 (0.069-0.097) | - |
| MMAF DAR 4 | 0.081 (0.067-0.098) | - |
| MMAE DAR 4 | 0.131 (0.115-0.150) | - |
| MMAE DAR 6 | 0.091 (0.078-0.106) | - |
| MMAE/F 2+2 | 0.129 (0.115-0.145) | - |
| MMAE/F 2+4 | 0.123 (0.110-0.138) | - |
| MMAE/F 4+2 | 0.149 (0.130-0.170) | - |

**Supplementary Table 2 $EC_{50}$ values in various cell lines (*n* = 4).** Values in parentheses are 95% confidential intervals.

| | $EC_{50}$ (nM) | | | | | | |
|---|---|---|---|---|---|---|---|
| | KPL-4 | JIMT-1 | JIMT-1 (MDR1+) | SKBR-3 | MDA-MB-231 | HepG2 | HEK293 |
| MMAF DAR 4 | 0.026 (0.023-0.0028) | 0.036 (0.032-0.041) | 0.012 (0.011-0.014) | 0.18 (0.13-0.29) | - | - | - |
| MMAE DAR 4 | 0.070 (0.062-0.078) | 0.064 (0.054-0.077) | - | 0.54 (0.38-0.83) | - | - | - |
| MMAE DAR 6 | 0.070 (0.057-0.094) | 0.060 (0.053-0.068) | - | Not tested | Not tested | Not tested | Not tested |
| MMAE/F 2+2 | 0.029 (0.025-0.033) | 0.045 (0.040-0.050) | Not tested | 0.34 (0.24-0.49) | - | - | - |
| MMAE/F 2+4 | 0.018 (0.016-0.020) | 0.026 (0.023-0.030) | 0.017 (0.015-0.019) | 0.23 (0.17-0.31 | - | - | - |
| MMAE/F 4+2 | 0.017 (0.015-0.019) | 0.024 (0.022-0.026) | 0.027 (0.022-0.032) | 0.18 (0.14-0.24) | - | - | - |

**Supplementary Table 3 Half-lives of N297A anti-HER2 mAb and ADCs at the elimination phase in mouse circulation (*n* = 4).**

| | $t_{1/2\beta\ \text{total mAb}}$ (day) | $t_{1/2\beta\ \text{ADC}}$ (day) |
|---|---|---|
| N297A anti-HER2 mAb | 17.4 | N/A |
| MMAF DAR 4 ADC | 13.3 | 14.7 |
| MMAE DAR 4 ADC | 14.6 | 15.1 |
| MMAE DAR 6 ADC | 14.2 | 15.7 |
| MMAE/F 2+2 ADC | 17.6 | 12.9 |
| MMAE/F 2+4 ADC | 14.9 | 12.2 |
| MMAE/F 4+2 ADC | 14.3 | 13.1 |

**Supplementary Table 4 Statistical significance.**

| Main Figures | Method | Comparison | Crude *P* value | Adjusted *P* value[a] | Significance |
|---|---|---|---|---|---|
| **Fig. 5b** | Welch's *t*-test | **Dose: 3 mg kg$^{-1}$ each** | | | |
| | | MMAE/F 4+2 *vs.* MMAE/F 2+4 (Day 38) | 0.0132 | 0.0396 | * |
| | | MMAE/F 4+2 *vs.* co-admin (Day 70) | 0.3590 | 0.7180 | n.s. |
| | | MMAE/F 4+2 *vs.* MMAE 4 (Day 87) | 0.3739 | 0.3739 | n.s. |
| **Fig. 5c** | Logrank (Mantel-Cox) | **Dose: 3 mg kg$^{-1}$ each** | | | |
| | | MMAE/F 4+2 *vs.* MMAE/F 2+4 | 0.0018 | 0.0054 | ** |
| | | MMAE/F 4+2 *vs.* co-admin | 0.0494 | 0.0988 | n.s. |
| | | MMAE/F 4+2 *vs.* MMAE 4 | 0.1343 | 0.1343 | n.s. |
| **Fig. 5d** | Welch's *t*-test | **Dose: 1 mg kg$^{-1}$ each** | | | |
| | | MMAE/F 4+2 *vs.* co-admin (Day 38) | 0.0002 | 0.0006 | *** |
| | | MMAE/F 4+2 *vs.* MMAE 4 (Day 38) | 0.0134 | 0.0268 | * |
| | | MMAE/F 4+2 *vs.* MMAE 6 (Day 66) | 0.0253 | 0.0253 | * |
| **Fig. 5e** | Logrank (Mantel-Cox) | **Dose: 1 mg kg$^{-1}$ each** | | | |
| | | MMAE/F 4+2 *vs.* co-admin | 0.0021 | 0.0063 | ** |
| | | MMAE/F 4+2 *vs.* MMAE 4 | 0.0024 | 0.0048 | *** |
| | | MMAE/F 4+2 *vs.* MMAE 6 | 0.0133 | 0.0133 | * |
| **Fig. 6d** | Welch's *t*-test | **Whole tumor (Cy5.5)** | | | |
| | | mAb-AF488/Cy5.5 *vs.* co-admin | 0.0292 | N/A | * |
| **Fig. 6e** | Welch's *t*-test | **Tumor section (AF488)** | | | |
| | | mAb-AF488/Cy5.5 *vs.* co-admin | 0.0057 | N/A | ** |

[a] Adjusted by the Holm–Bonferroni method. *$P < 0.05$; **$P < 0.01$; ***$P < 0.005$; n.s., not significant.

#### 2. SUPPLEMENTARY NOTES

##### General information

Unless otherwise noted, all materials for chemical synthesis were purchased from commercial suppliers (for example, Acros Organics, AnaSpec, Broadpharm, Chem-Impex International, Thermo Fisher Scientific, Levena Biopharma, Millipore Sigma, and TCI America) and used as received. All anhydrous solvents were purchased and stored over activated molecular sieves under argon atmosphere.

Nuclear magnetic resonance (NMR) spectra were recorded using a Bruker DPX spectrometer ($^{1}H$: 300 MHz, $^{13}C$: 75 MHz) using chloroform-*d* ($CDCl_3$), methanol-$d_4$ ($CD_3OD$), or dimethyl sulfoxide-$d_6$ (DMSO-$d_6$) as deuterated solvent. Chemical shifts (δ) in $^{1}H$ and $^{13}C$ NMR spectra were reported in parts per million (ppm) relative to $CDCl_3$ ($^{1}H$: δ = 7.26 ppm, $^{13}C$: δ = 77.16 ppm), $CD_3OD$ ($^{1}H$: δ = 3.33 ppm, $^{13}C$: δ = 49.0 ppm) or DMSO-$d_6$ ($^{13}C$: δ = 39.52 ppm). Coupling constants (*J*) in all NMR spectra are reported in Hertz (Hz).

Analytical reverse-phase high performance liquid chromatography (RP-HPLC) was performed using an Agilent LC-MS system consisting of a 1100 HPLC and a 1946D single quadrupole electrospray ionization (ESI) mass spectrometer equipped with a C18 reverse-phase column (small molecules: Accucore C18 column, 3×50 mm, 2.6 µm, Thermo Scientific; antibodies: MabPac RP column, 3×50 mm, 4 µm, Thermo Scientific). Standard analysis conditions for organic molecules were as follows: flow rate = 0.5 mL/min; solvent A = water containing 0.1% formic acid; solvent B = acetonitrile containing 0.1% formic acid. Compounds were analyzed using a linear gradient and monitored with UV detection at 210 and 254 nm. Preparative HPLC was performed using a Breeze HPLC system (Waters) equipped with a C18 reverse-phase column (SunFire Prep C18 OBD, 19×150 mm, 5.0 µm; Waters). Standard purification conditions were as follows: flow rate = 20 mL/min; solvent A = water containing 0.05% trifluoroacetic acid (TFA), 0.1% formic acid or 0.1% $NH_4OH$; solvent B = acetonitrile containing 0.05% TFA, 0.1% formic acid, or 0.1% $NH_4OH$. Compounds were analyzed using a linear gradient and monitored with UV detection at 210 and 254 nm. In all cases, fractions were analyzed off-line using the LC-MS for purity confirmation and those containing a desired product were lyophilized using a Labconco Freezone 4.5 Liter Benchtop Freeze Dry System. High-resolution mass spectra were obtained using an Agilent 6530 Accurate Mass Q-TOF LC/MS at the UT Austin Mass Spectrometry Core Facility.

#### Synthesis

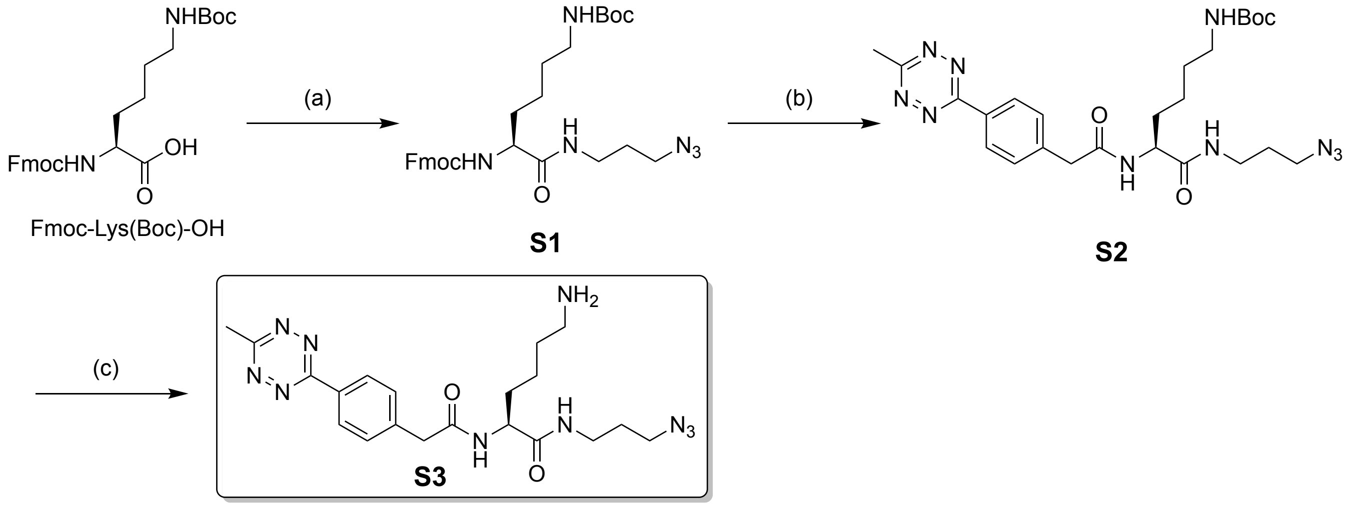

Synthesis of azido-methyltetrazine di-arm linker **S3**. Reagents and conditions: (a) 3-azidopropylamine, EDC·HCl, NHS, DMF, room temp, overnight; (b) 30% piperidine/DMF, room temp, 1 h; then methyltetrazine acid, EDC·HCl, NHS, DMF, room temp, 2 h; (c) 20% TFA/DCM, room temp, 1 h.

**Fmoc-Lys(Boc)-$N_3$ (S1).** Fmoc-Lys(Boc)-OH (234.3 mg, 0.50 mmol) in DMF (3 mL) was mixed with *N*-hydroxysuccinimide (NHS, 115.1 mg, 1.0 mmol) and *N*-(3-dimethylaminopropyl)-*N'*-ethylcarbodiimide hydrochloride (EDC·HCl, 191.7 mg, 1.0 mmol) at room temperature. To the solution was added 3-azidopropylamine (65 μL, 0.65 mmol) and the resulting mixture was stirred overnight at room temperature. The reaction was quenched with 15% citric acid and extracted with ethyl acetate. The organic layer was washed with brine, dried over $Na_2SO_4$, and concentrated in vacuo. The residue was purified by column chromatography using a Biotage Isolera Flash Purification System (0–20% of DCM/MeOH, 20 mL/min, SNAP cartridge KP-Sil 10 g) to afford **S1** (431.9 mg, quant.). White solid. $^1$H NMR (300 MHz, $CDCl_3$) δ 7.77 (d, *J* = 7.5 Hz, 2H), 7.59 (d, *J* = 7.4 Hz, 2H), 7.41 (t, *J* = 7.2 Hz, 2H), 7.31 (td, *J* = 7.4, 1.2 Hz, 2H), 6.39 (brs, 1H), 4.63 (brs, 1H), 4.41 (d, *J* = 6.9 Hz, 2H), 4.21 (t, *J* = 6.8 Hz, 1H), 4.09 (brs, 1H), 3.34 (t, *J* = 6.4 Hz, 4H), 3.14–3.08 (m, 1H), 2.06–1.69 (m, 6H), 1.56–1.47 (m, 2H), 1.43 (s, 9H), 1.40–1.31 (m, 2H). $^{13}$C NMR (75 MHz, $CDCl_3$) δ 172.1, 156.50, 156.48, 143.89, 143.86, 141.5 (2 carbons), 127.9 (2 carbons), 127.2 (2 carbons), 125.2 (2 carbons), 120.2 (2 carbons), 79.5, 67.2, 55.1, 49.4, 47.3, 39.8, 37.3, 32.0, 29.7, 28.8, 28.6 (3 carbons), 22.6. HRMS (ESI) Calcd. For $C_{29}H_{38}N_6O_5Na$ $[M+Na]^+$: 573.2796. Found: 573.2801.

**Methyltetrazine-Lys(Boc)-$N_3$ (S2).** Compound **S1** (69.0 mg, 0.13 mmol) was dissolved in DMF (350 μL) and then piperidine (150 μL) was added to the solution at room temperature. After 1 h, the mixture was concentrated in vacuo and the crude products were precipitated with cold diethyl ether (5–6 mL) followed by centrifugation at 2,000 × *g* for 3 min (3 times). The pellet was dried in vacuo and used immediately in the next step without purification. To the residue were added methyltetrazine acid (34.5 mg, 0.15 mmol), NHS (26.5 mg, 0.25 mmol), and EDC·HCl (47.8 mg, 0.25 mmol) in DMF (1 mL). After being stirred at room temperature for 2 h, the reaction mixture was quenched with 15% citric acid and extracted with ethyl acetate. The organic layer was washed with brine, dried over $Na_2SO_4$, and concentrated. The residue was purified by preparative RP-HPLC under acidic conditions to afford **S2** (10.0 mg, 14% for the 2 steps). Red powder. $^1$H NMR (300 MHz, $CDCl_3$) δ 8.53 (d, *J* = 8.3 Hz, 2H), 7.47 (d, *J* = 8.1 Hz, 2H), 4.68 (t, *J* = 5.1 Hz, 1H), 4.43–4.36 (m, 1H), 3.65 (s, 2H), 3.34–3.22 (m, 4H), 3.09 (s, 3H), 3.07–3.01 (m, 1H), 1.90–1.58 (m, 6H), 1.53–1.45 (m, 2H), 1.43 (s, 9H), 1.37–1.25 (m, 2H). $^{13}$C NMR (75 MHz, $CD_3OD$) δ 174.4, 173.2, 168.7, 165.2, 158.5, 141.8, 132.2, 131.2 (2 carbons), 129.0 (2 carbons), 79.9, 55.1, 50.0, 43.4, 41.1, 37.7, 32.8, 30.5, 29.6, 28.8 (3 carbons), 24.2, 21.0. HRMS (ESI) Calcd. For $C_{25}H_{36}N_{10}O_4Na$ $[M+Na]^+$: 563.2813. Found: 563.2822.

**Azido-methyltetrazine di-arm linker (S3).** Compound **S2** (10.0 mg, 18 μmol) was dissolved in DCM (240 μL) and TFA (60 μL) was added to the solution at room temperature. After 1 h, the mixture was concentrated and purified by preparative RP-HPLC under acidic conditions to afford di-arm linker **S3** (3.3 mg, 42%). Red powder. $^1$H NMR (300 MHz, $CD_3OD$) δ 8.53 (d, *J* = 8.4 Hz, 2H), 8.42 (d, *J* = 7.7 Hz, 1H), 8.08 (t, *J* = 5.6 Hz, 1H), 7.60 (d, *J* = 8.3 Hz, 2H), 4.40–4.26 (m, 1H), 3.73 (s, 2H), 3.39–3.34 (m, 2H), 3.31–3.23 (m, 2H), 3.06 (s, 3H), 2.89 (t, *J* = 7.1 Hz, 2H), 1.84–1.59 (m, 6H), 1.50–1.35 (m, 2H). $^{13}$C NMR (75 MHz, DMSO-$d_6$) δ 171.5, 169.6, 167.1, 163.3, 141.3, 130.2 (2 carbons), 130.1, 127.3 (2 carbons), 52.5, 48.3 (2 carbons), 42.0, 35.9, 31.6, 28.3, 26.7, 22.4, 20.9. HRMS (ESI) Calcd. For $C_{20}H_{29}N_{10}O_2$ $[M+H]^+$: 441.2469. Found: 441.2460. This linker was used to prepare the MMAE/F 2+2 ADC, the MMAE/F 2+4 ADC, the DAR 4 single-drug ADCs, and the single- and dual-dye conjugates.

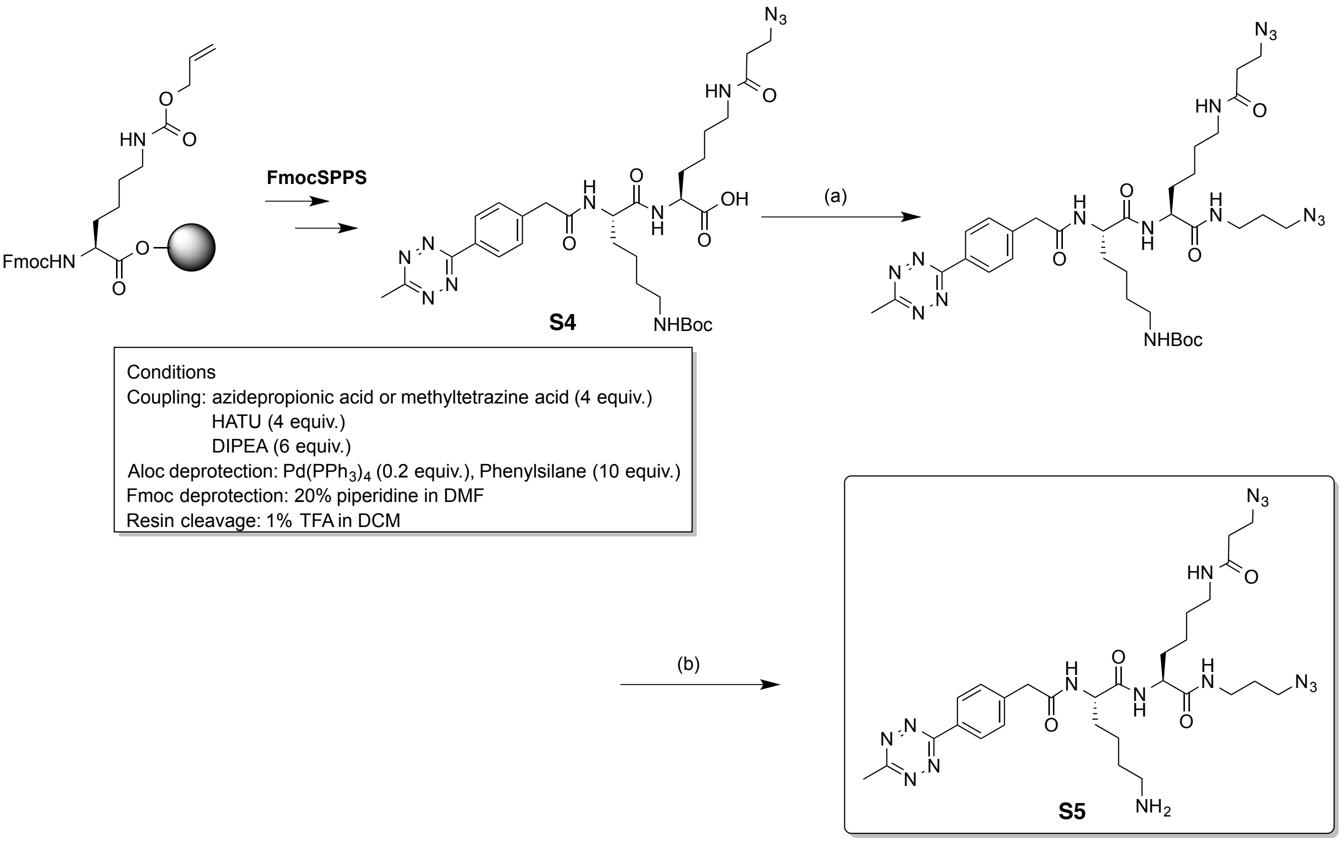

Synthesis of diazido-methyltetrazine tri-arm linker (**S5**). Reagents and conditions: (a) 3-azidopropylamine, EDC·HCl, NHS, DMF, room temp, 2 h; (b) 20% TFA/DCM, room temp, 1 h.

**Synthesis of MeTet-Lys(Boc)-Lys($N_3$)-OH (S4).** 2-Chlorotrityl chloride resin (500 mg, 0.8 mmol, AnaSpec) and Fmoc-Lys(Alloc)-OH (362 mg, 0.8 mmol) were taken to a manual solid-phase reactor containing *N*,*N*-diisopropylethylamine (DIPEA, 209 µL, 1.2 mmol) and DMF (3 mL) and agitated for 2 h. MeOH (600 µL, 1.5 mmol) was added to the resin and agitated for 20 min. The solution was drained and the resin was washed with DMF (5×3 mL) and DCM (5×3 mL). To remove the Alloc-protecting group before initial coupling, the resin was treated with $Pd(PPh_3)_4$ (138.7 mg, 0.12 mmol) and phenylsilane (49.3 µL, 0.4 mmol) in DCM (3 mL) under Ar atmosphere (15 min x 2) and washed with DMF (5×3 mL) and DCM (5×3 mL). To remove a Fmoc-protecting group after each coupling, the resin was treated with 20% piperidine/DMF (5 mL) for 20 min and washed with DMF (5×3 mL) and DCM (5×3 mL). Azidopropionic acid, Fmoc-Lys(Boc)-OH, or methyltetrazine acid (2 equiv.) was pre-activated using 1-[bis(dimethylamino)methylene]-1H-1,2,3-triazolo[4,5-b]pyridinium 3-oxid hexafluorophosphate (HATU, 2 equiv.) and DIPEA (3 equiv.) in DMF for 5 min, and the cocktail was added to the resin. The resin was agitated at room temperature for 1.5 h. Completion of the coupling was verified by the Kaiser test. After each coupling step, the coupling cocktail was drained and the resin was washed with DMF (5×3 mL) and DCM (5×3 mL). The resulting protected peptide resin was treated with a cocktail of 1% trifluoroacetic acid (TFA)/DCM at room temperature for 1 h. The solution was concentrated in vacuo and the crude peptide was precipitated with cold diethyl ether (5–6 mL) followed by centrifugation at 2,000 × g for 3 min (3 times). The resulting crude products were dried in vacuo and purified by RP-HPLC to afford analytically pure peptide **S4** (88.3 mg, 31% based on the resin loading rate). Red powder. $^1$H NMR (300 MHz, $CD_3OD$) δ 8.49 (d, *J* = 8.3 Hz, 2H), 7.56 (d, *J* = 8.3 Hz, 2H), 4.41–4.32 (m, 2H), 3.71 (s, 2H), 3.50 (t, *J* = 6.5 Hz, 2H), 3.21–3.05 (m, 2H), 3.03 (s, 3H), 3.03–2.96 (m, 2H), 2.38 (t, *J* = 6.2 Hz, 2H), 1.92–1.77 (m, 2H), 1.77–1.61 (m, 2H), 1.56–1.44 (m, 6H), 1.43 (s, 9H), 1.41–1.35 (m, 2H). HRMS (ESI) Calcd. For $C_{31}H_{45}N_{11}O_7Na$ $[M+Na]^+$: 706.3396. Found: 706.3396.

**Synthesis of diazido-methyltetrazine tri-arm linker (S5).** To compound **S4** (12.6 mg, 0.018 mmol) was added a DMF solution (1 mL) of 3-azidopropylamine (2.4 µL, 0.023 mmol), NHS (7.0 mg, 0.036 mmol), and EDC·HCl (4.2 mg, 0.036 mmol). After being stirred at room temperature for 2 h, the reaction mixture was quenched with 15% citric acid and extracted with ethyl acetate. The organic layer was washed with brine, dried over $Na_2SO_4$, and concentrated and used immediately in the next step without purification. To the residue DCM (0.2 mL) and TFA (0.2 mL) were added at room temperature. After 1 h, the mixture was concentrated and purified by preparative RP-HPLC under acidic conditions to afford tri-arm linker **S5** (4.0 mg, 33 %). Red powder. $^1$H NMR (300 MHz, $CD_3OD$) δ 8.50 (d, *J* = 8.3 Hz, 2H), 7.57 (d, *J* = 8.3 Hz, 2H), 4.41–4.29 (m, 1H), 4.28–4.17 (m, 1H), 3.72 (s, 2H), 3.52 (t, *J* = 6.4 Hz, 2H), 3.39–3.32 (m, 2H), 3.29–3.06 (m, 4H), 3.04 (s, 3H), 2.92 (t, *J* = 7.5 Hz, 2H), 2.40 (t, *J* = 6.4 Hz, 2H), 1.91–1.58 (m, 8H), 1.54–1.26 (m, 6H). HRMS (ESI) Calcd. For $C_{29}H_{43}N_{15}O_4Na$ $[M+Na]^+$: 688.3515. Found: 688.3538. This linker was used to prepare MMAE/F 4+2 and DAR 6 ADCs.

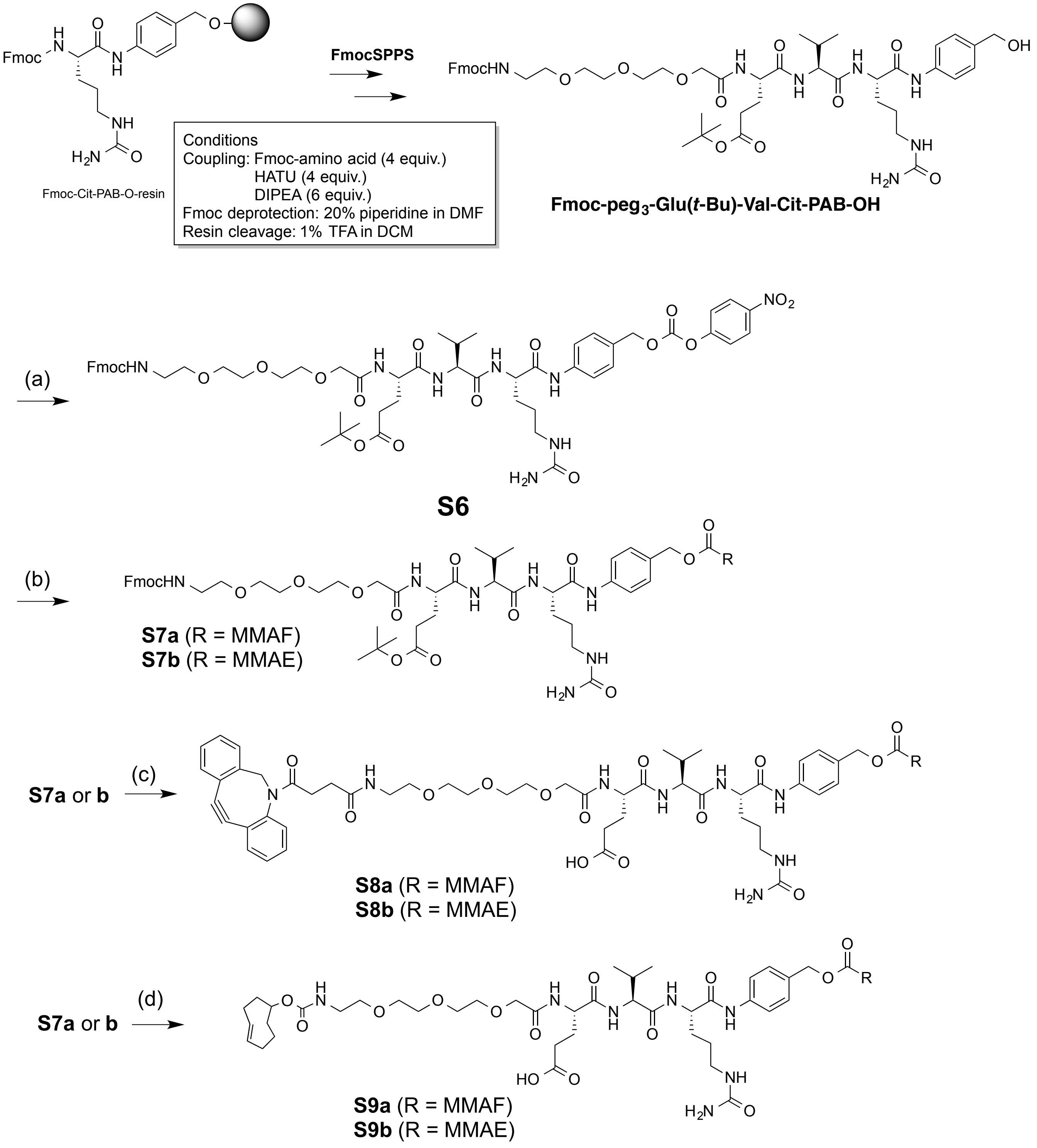

Synthesis of **S8a,b** and **S9a,b.** Reagents and conditions: (a) bis(4-nitrophenyl) carbonate, DMAP, DMF, room temp, 2 h; (b) MMAF (for **S7a**) or MMAE (for **S7b**), DIPEA, HOAt, DMF, 37 °C, overnight.; (c) 50% TFA/DCM, room temp, 1 h; 50% diethylamine/DMF, room temp, 1 h, then DBCO-NHS; (d) 50% TFA/DCM, room temp, 1 h; 50% diethylamine/DMF, room temp, 1 h, then TCO-NHS, DIPEA, room temp, overnight.

**Synthesis of Fmoc-peg$_3$-Glu(*t*-Bu)-Val-Cit-PAB-OH.** Fmoc-Cit-PAB-O-resin was prepared according to the procedure described previously[1]. To remove the Fmoc-protecting group after each coupling, the resin was treated with 20% piperidine/DMF (5 mL) for 20 min and washed with DMF (5×3 mL) and DCM (5×3 mL). Fmoc-protected amino acid (2 equiv.) was pre-activated using HATU (2 equiv.) and DIPEA (3 equiv.) in DMF for 3 min, and the cocktail was added to the resin. The resin was agitated at room temperature for 1 h. The completion of the coupling was verified by the Kaiser test. After each coupling step, the coupling cocktail was drained and the resin was washed with DMF (5×3 mL) and DCM (5×3 mL). The resulting protected peptide resin was treated with cocktail of 1% trifluoroacetic acid (TFA)/DCM at room temperature for 1 h. The solution was concentrated in vacuo and the resulting crude products were precipitated with cold diethyl ether (5–6 mL) followed by centrifugation at 2,000 × g for 3 min (3 times). The peptide pellet was dried in vacuo and used immediately in the next step without purification.

**Synthesis of Fmoc-peg$_3$-Glu(*t*-Bu)-Val-Cit-PABC-PNP (S6).** Bis(2,4-dinitrophenyl) carbonate (154 mg, 0.51 mmol) and DMAP (24.9 mg, 0.20 mmol) were added to a crude mixture of Fmoc-peg$_3$-Glu(*t*-Bu)-Val-Cit-PAB-OH (97.6 mg, assumed to be 0.10 mmol) in DMF (4 mL), and the mixture was stirred at room temperature for 2 h. The crude products were purified by preparative RP-HPLC to give analytically pure peptide **S6** (68.2 mg, 59% overall yield, calculated based on the resin loading rate). Purity was confirmed by LC-MS. Off-white powder. HRMS (ESI) Calcd. For $C_{57}H_{72}N_8O_{17}Na$ $[M+Na]^+$: 1163.4908. Found: 1163.4911.

**Synthesis of Fmoc-peg$_3$-Glu(*t*-Bu)-Val-Cit-PABC-MMAF (S7a).** A solution of **S6** (7.0 mg, 6.1 μmol) in DMF (620 μL) was mixed with monomethyl auristatin F TFA salt (6.7 mg, 8.0 μmol), HOAt (1.7 mg, 0.012 mmol), and DIPEA (5.3 μL, 0.031 mmol). The resulting mixture was stirred overnight at 37 °C. The crude products were directly purified by preparative RP-HPLC to afford peptide **S7a** (7.8 mg, 73%). Purity was confirmed by LC-MS. White powder. HRMS (ESI) Calcd. $C_{90}H_{132}N_{12}O_{22}Na_2$ $[M+2Na]^{2+}$: 889.4682. Found: 889.4695. Peptide **S7b** was synthesized in a similar manner.

**Fmoc-peg$_3$-Glu(*t*-Bu)-Val-Cit-PABC-MMAE (S7b).** Monomethyl auristatin E TFA salt was used instead of monomethyl auristatin F TFA salt. Yield: 7.7 mg, 84%. White powder. HRMS (ESI) Calcd. For $C_{90}H_{134}N_{12}O_{21}Na_2$ $[M+2Na]^{2+}$: 882.4785. Found: 882.4817.

**DBCO-peg$_3$-Glu-Val-Cit-PABC-MMAF (S8a).** TFA (500 μL) was added to a DCM solution (500 μL) of compound **S7a** (6.2 mg, 3.6 μmol) at 0 ºC. After being stirred at room temperature for 1 h, the reaction mixture was concentrated in vacuo and the crude peptide was precipitated with cold diethyl ether (5–6 mL) followed by centrifugation at 2,000 × g for 3 min (3 times). The resulting crude products were dried in vacuo. Diethylamine (300 μL) was added to the residue dissolved in DMF (300 μL), and the mixture was stirred at room temperature for 1 h. The solution was concentrated and dried in vacuo and used in the next step without further purification. DBCO-NHS (1.5 mg, 5.4 μmol) was added to a solution of this crude mixture in DMF (500 μL) and the mixture was stirred overnight at room temperature. The crude products were purified by preparative RP-HPLC under basic conditions to afford peptide **S8a** (4.2 mg, 67% for the 3 steps). Purity was confirmed by LC-MS. This compound has previously been characterized by us[1]. White powder. Peptide **S8b** was prepared from **S7b** in a similar manner.

**DBCO-peg$_3$-Glu-Val-Cit-PABC-MMAE (S8b).** Yield: 3.0 mg, 41% for the 3 steps. White powder. HRMS (ESI) Calcd. For $C_{90}H_{129}N_{13}O_{21}Na_2$ $[M+2Na]^{2+}$: 886.9605. Found: 886.9629.

**TCO-peg$_3$-Glu-Val-Cit-PABC-MMAF (S9a).** TFA (300 μL) was added to a DCM solution (300 μL) of compound **S7a** (7.8 mg, 4.5 μmol). After being stirred at room temperature for 1 h, the reaction mixture was concentrated in vacuo and the crude peptide was precipitated with cold diethyl ether (5–6 mL) followed by centrifugation at 2,000 × g for 3 min (3 times). The resulting crude products were dried in vacuo and used in the next step without further purification. Diethylamine (250 μL) was added to a half of this crude mixture in DMF (250 μL), and the mixture was stirred at room temperature for 1 h. The solution was concentrated and dried in vacuo and used in the next step without further purification. To a solution of this crude mixture in DMF (300 μL) was added TCO-NHS (0.8 mg, 2.9 μmol) and DIPEA (0.8 μL, 4.5 μmol), and the mixture was stirred at room temperature for 3 h. The crude products were purified by preparative RP-HPLC under basic conditions to afford peptide **S9a** (2.0 mg, 54% for the 3 steps). Purity was confirmed by LC-MS. White powder. HRMS (ESI) Calcd. For $C_{80}H_{126}N_{12}O_{22}Na_2$ $[M+2Na]^{2+}$: 826.4447. Found: 826.4461. Peptide **S9b** was prepared from peptide **S7b** in a similar manner.

**TCO-peg$_3$-Glu-Val-Cit-PABC-MMAE (S9b).** Yield: 4.8 mg, 50% for the 3 steps. White powder. HRMS (ESI) Calcd. For $C_{80}H_{128}N_{12}O_{21}Na_2$ $[M+2Na]^{2+}$: 819.4551. Found: 819.4560.

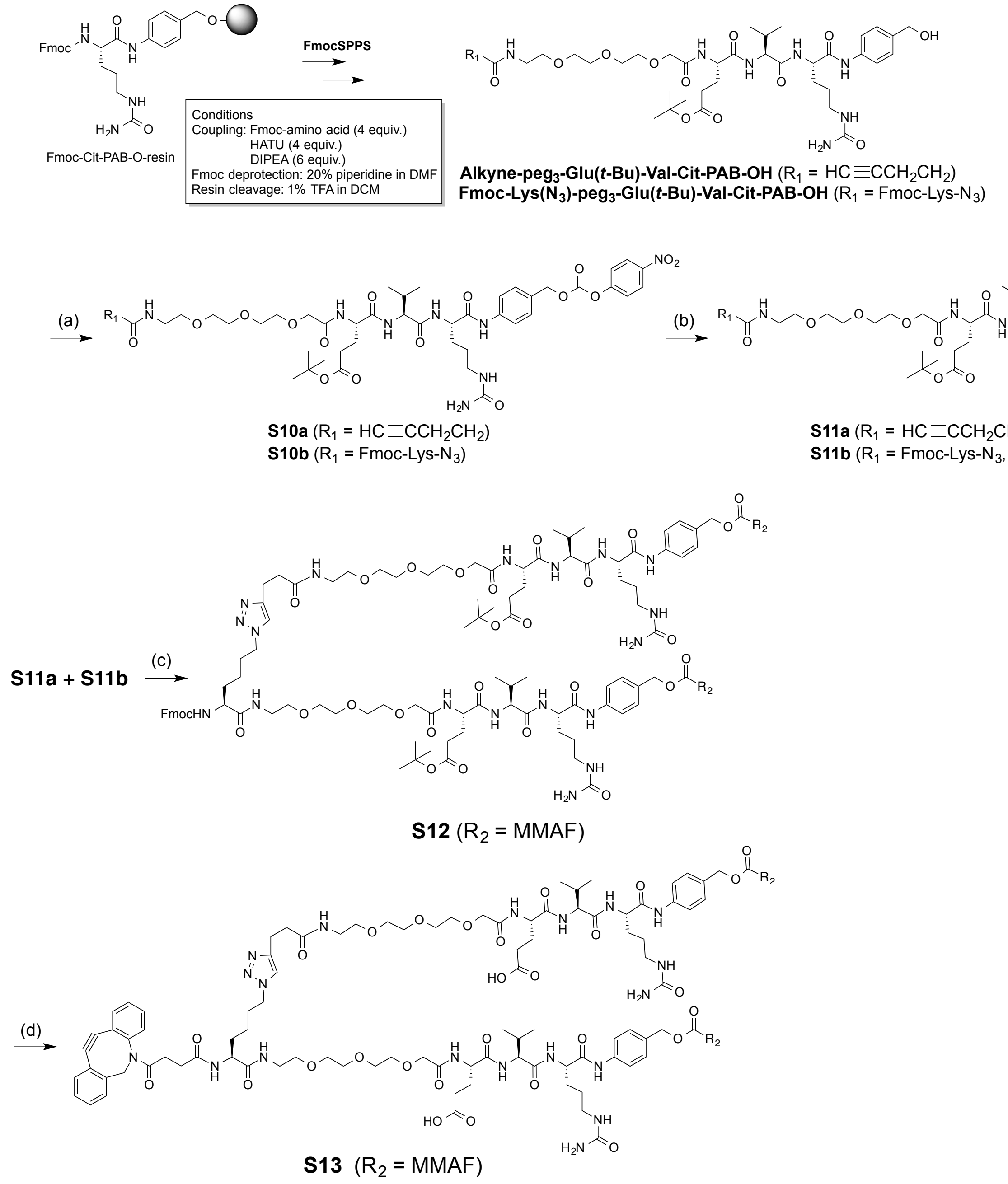

Synthesis of di-MMAF module **S13**. Reagents and conditions: (a) bis(4-nitrophenyl) carbonate, DMAP, DMF, room temp, 2 h; (b) MMAF, DIPEA, HOAt, DMF, 37 °C, overnight; (c) $CuSO_4 \cdot 5H_2O$, TBTA, sodium ascorbate, 1M phosphate buffer pH 7.0, DMSO, room temp, 2 h; (d) 20% TFA/DCM, room temp, 1 h; 50% diethylamine/DMF, room temp, 2 h; then DBCO-NHS, DIPEA, room temp, overnight.

**Alkyne-peg$_3$-Glu(*t*-Bu)-Val-Cit-PABC-PNP (S10a).** Crude alkyne-peg$_3$-Glu(*t*-Bu)-Val-Cit-PAB-OH was prepared in a similar manner as described earlier. This crude peptide (53.0 mg, assumed to be 0.064 mmol) was converted into peptide **S10a** in the same manner as described in the synthesis of **S6**. Yield:

10.4 mg, 61% overall yield based on the resin loading rate. White powder. HRMS (ESI) Calcd. For $C_{47}H_{66}N_8O_{16}Na$ $[M+Na]^+$: 1021.4489. Found: 1021.4495.

**Fmoc-Lys($N_3$)-peg$_3$-Glu(*t*-Bu)-Val-Cit-PABC-PNP (S10b).** Crude Fmoc-Lys($N_3$)-peg$_3$-Glu(*t*-Bu)-Val-Cit-PAB-OH was prepared in a similar manner as described earlier. This crude peptide (43.6 mg, assumed to be 0.039 mmol) was converted into peptide **S10b** in the same manner as described in the synthesis of **S6**. Yield: 10.7 mg, 45% overall yield based on the resin loading rate. Purity was confirmed by LC-MS. White powder. HRMS (ESI) Calcd. For $C_{63}H_{82}N_{12}O_{18}Na$ $[M+Na]^+$: 1317.5762. Found: 1317.5758.

**Alkyne-peg$_3$-Glu(*t*-Bu)-Val-Cit-PABC-MMAF (S11a).** Prepared from **S10a** in the same manner as the synthesis of **S7a.** Yield: 9.1 mg, 69%. White powder. HRMS (ESI) Calcd. For $C_{80}H_{126}N_{12}O_{21}Na_2$ $[M+2Na]^{2+}$: 818.4472. Found: 818.4492.

**Fmoc-Lys($N_3$)-peg$_3$-Glu(*t*-Bu)-Val-Cit-PABC-MMAF (S11b).** Prepared from **S10b** in the same manner as the synthesis of **S7a.** Yield: 6.6 mg, 42%. White powder. HRMS (ESI) Calcd. For $C_{96}H_{142}N_{16}O_{23}Na_2$ $[M+2Na]^{2+}$: 966.5109. Found: 966.5140.

**Fmoc-Lys-(peg$_3$-Glu(*t*-Bu)-Val-Cit-PABC-MMAF)$_2$ (S12).** A mixture of alkyne peptide **S11a** (200 μL, 10 mM in DMSO), azide peptide **S11b** (100 μL, 10 mM in DMSO), and 1 M phosphate buffer (pH 7.0, 300 μL) was treated with a mixture of $CuSO_4 \cdot 5H_2O$ (200 μL, 20 mM in DI water) and tris[(1-benzyl-1H-1,2,3-triazol-4-yl)methyl]amine (TBTA, 200 μL, 40 mM in DMSO). DMSO (900 μL) and sodium ascorbate (100 μL, 100 mM in deionized water) was added to the mixture and stirred at 37 °C for 2 h. The crude products were directly purified by preparative RP-HPLC to give analytically pure peptide **S12** (5.1 mg, 73%). Purity was confirmed by LC-MS. Yellow powder. HRMS (ESI) Calcd. For $C_{176}H_{270}N_{28}O_{44}$ $[M+2H]^{2+}$:1739.9870. Found: 1739.9913.

**DBCO-Lys-(peg$_3$-Glu-Val-Cit-PABC-MMAF)$_2$ (S13).** TFA (60 μL) was added to a DCM solution (240 μL) of compound **S12** (5.1 mg, 1.5 μmol). After being stirred at room temperature for 1 h, the reaction mixture was concentrated in vacuo and the crude products were precipitated with cold diethyl ether (5–6 mL) followed by centrifugation at 2,000 × g for 3 min (3 times). The resulting crude products were dried in vacuo. To the residue in DMF (200 μL) was added diethylamine (200 μL), and the mixture was stirred at room temperature for 2 h. The solution was concentrated and dried in vacuo and used in the next step without further purification. To a solution of this crude mixture in DMF (0.2 mL) was added DBCO-NHS (1.5 mg, 3.7 µmol) and DIPEA (1 µL, 5.7 µmol), and the mixture was stirred overnight at room temperature. The crude products were directly purified by preparative RP-HPLC under basic conditions to afford peptide **S13** (0.5 mg, 10% for the 3 steps). Purity was confirmed by LC-MS. White powder. HRMS (ESI) Calcd. For $C_{172}H_{255}N_{29}O_{44}Na_2$ $[M+2Na]^{2+}$: 1738.4196. Found: 1738.4167.

**Expression and purification of human monoclonal antibodies**. All humanized mAbs were produced according to the procedure reported previously[2,3]. Free style HEK-293 human embryonic kidney cells (Invitrogen) were transfected with a mammalian expression vector encoding for the human IgG1 kappa light chain and full length heavy chain sequences (based on the variable sequences of trastuzumab and sacituzumab). A mutation of N297A was incorporated into the heavy chain constant region to produce aglycosylated mAbs. The transfected HEK-293 cells were cultured in a humidified cell culture incubator at 37 °C with 8% $CO^2$ and shaking at 150 rpm for 7 days before harvesting the culture medium. The antibody secreted into the culture medium was purified using Protein A resin (GE Healthcare).

**Flow cytometry.** DAPI was added into all samples for live cell selection. First, unstained cells were used to set up the scatter, single cell selection, and anti-HER2-FITC positive area. Next, IgG-FITC treated cells were used to confirm the specificity of anti-HER2-FITC. After confirming no non-specific binding of IgG-FITC, anti-HER2-FITC treated cells were applied for measuring the HER2-positive cells. Total 10,000 cells were counted from each sample.

### NMR, HPLC, and deconvoluted ESI-MS Data

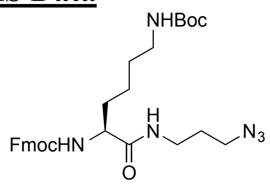

**S1**

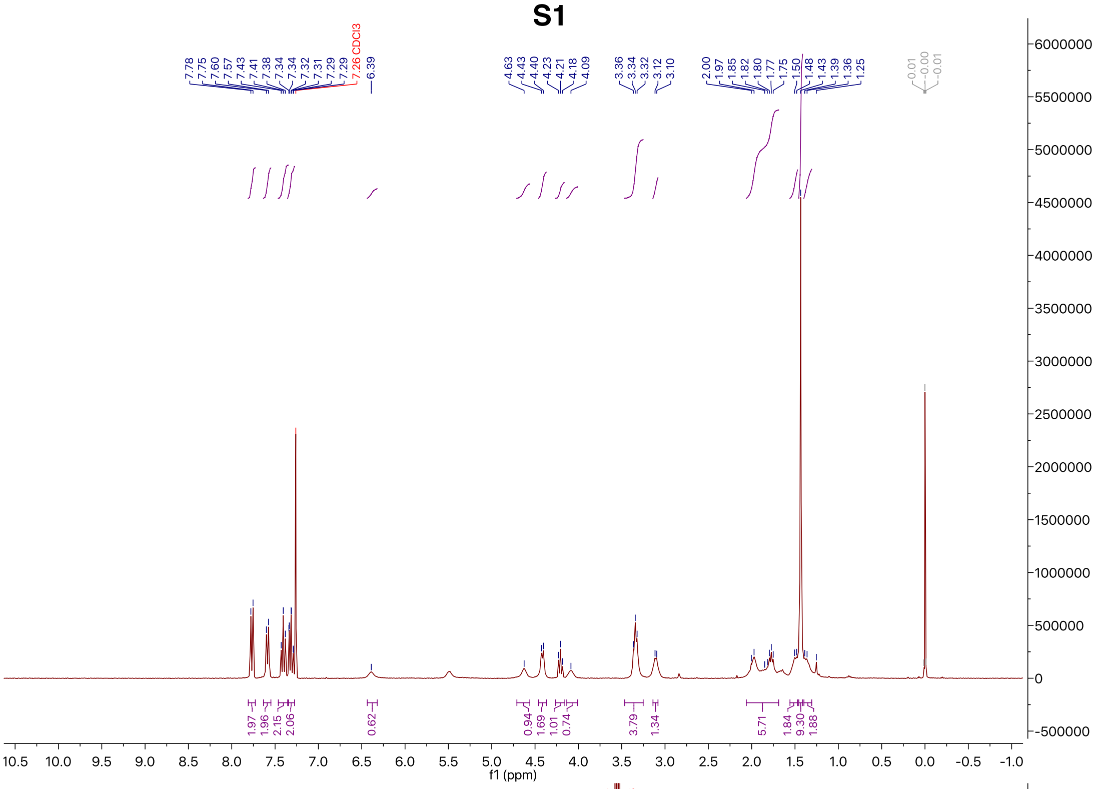

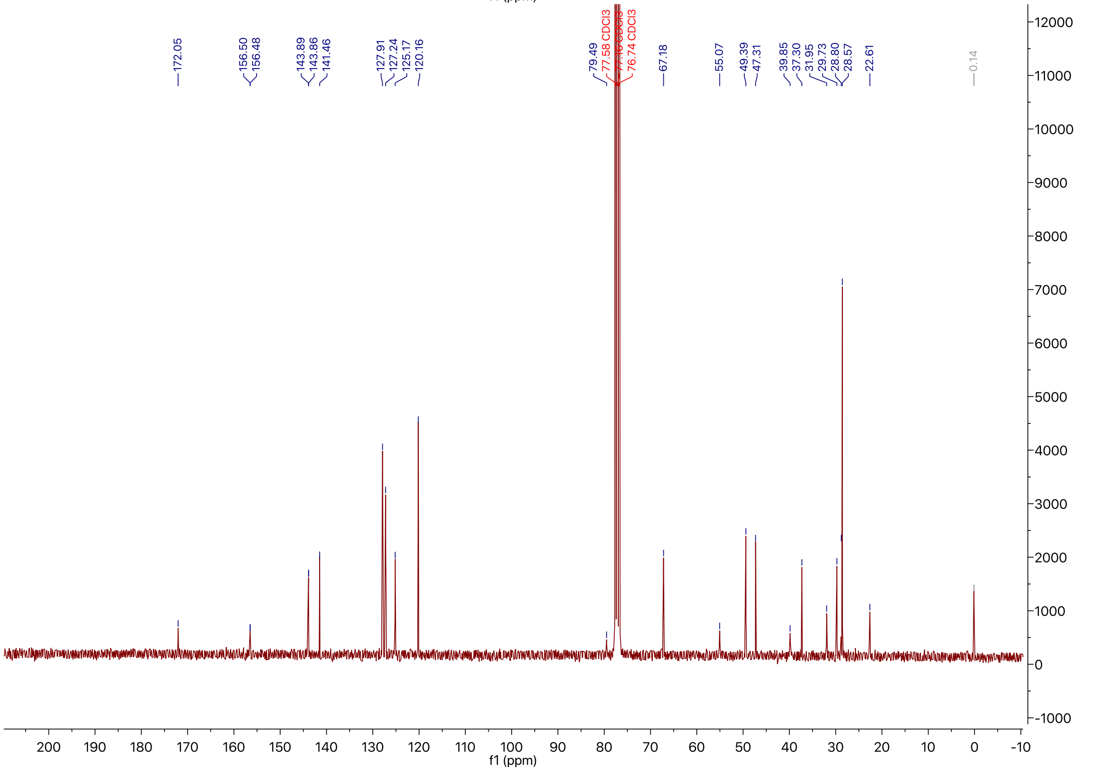

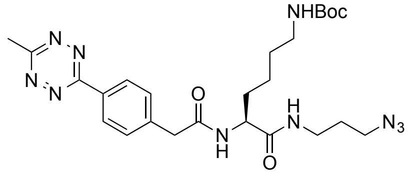
NHBoc
N
N
N
N
O
N
H
H
N
N3
O

**S2**

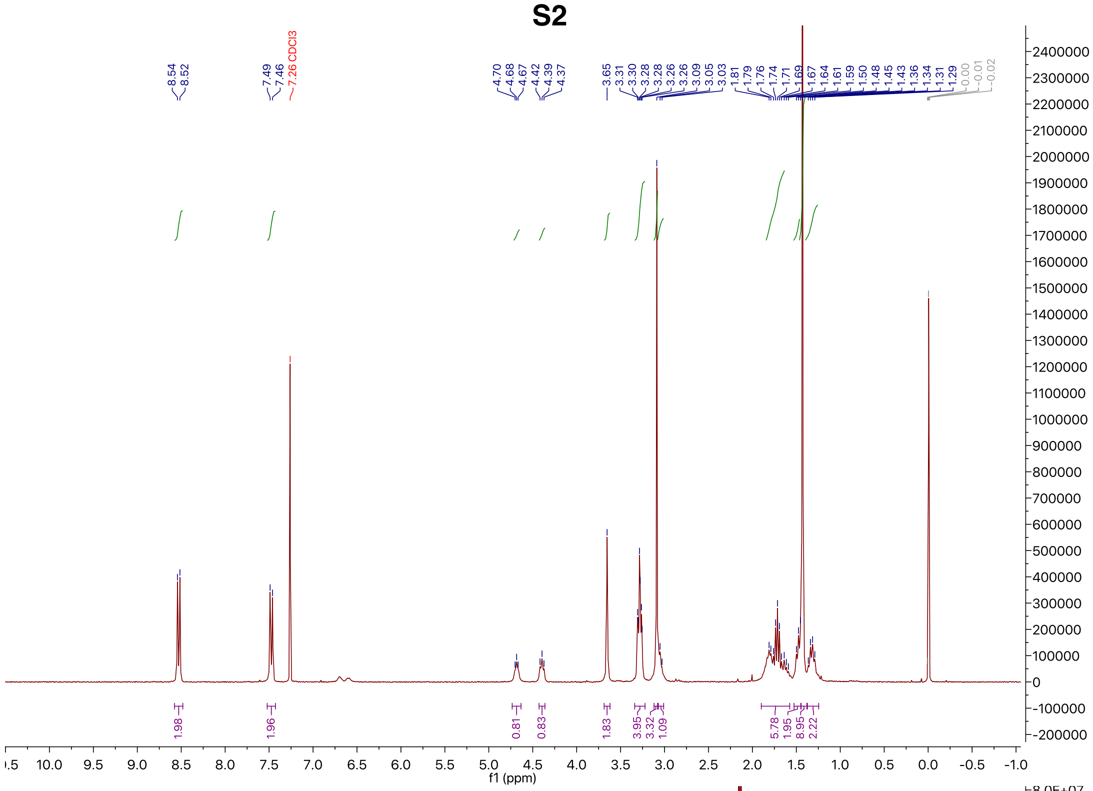
8.54
8.52
7.49
7.46
7.26 CDCl3
4.70
4.68
4.67
4.42
4.39
4.37
3.65
3.31
3.30
3.28
3.28
3.26
3.26
3.09
3.05
3.03
1.81
1.79
1.76
1.74
1.71
1.69
1.67
1.64
1.61
1.59
1.50
1.48
1.45
1.43
1.36
1.34
1.31
1.29
0.00
-0.01
-0.02
1.98
1.96
0.81
0.83
1.83
3.95
3.32
1.09
5.78
1.95
8.95
2.22
f1 (ppm)

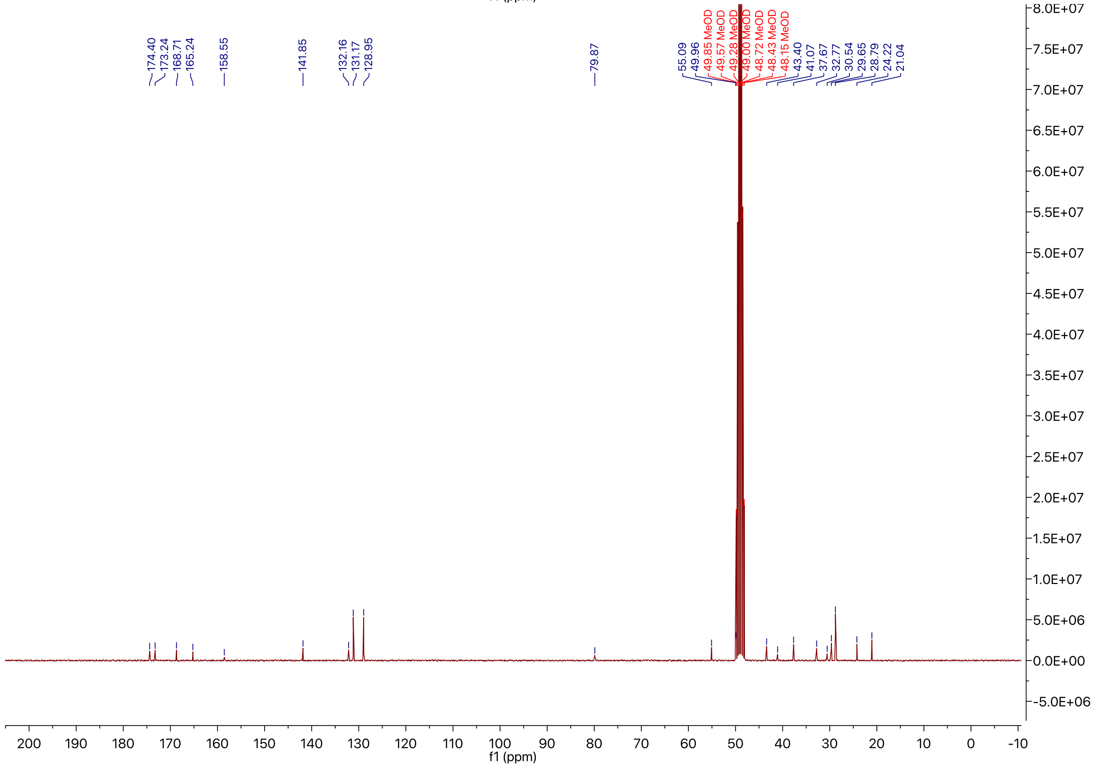
174.40
173.24
168.71
165.24
158.55
141.85
132.16
131.17
128.95
79.87
55.09
49.96
49.85 MeOD
49.57 MeOD
49.28 MeOD
49.00 MeOD
48.72 MeOD
48.43 MeOD
48.15 MeOD
43.40
41.07
37.67
32.77
30.54
29.65
28.79
24.22
21.04
f1 (ppm)

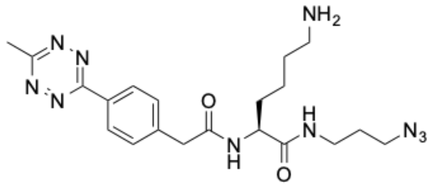
NH2
N
N
N
N
O
N
H
H
N
O
N3

**S3**

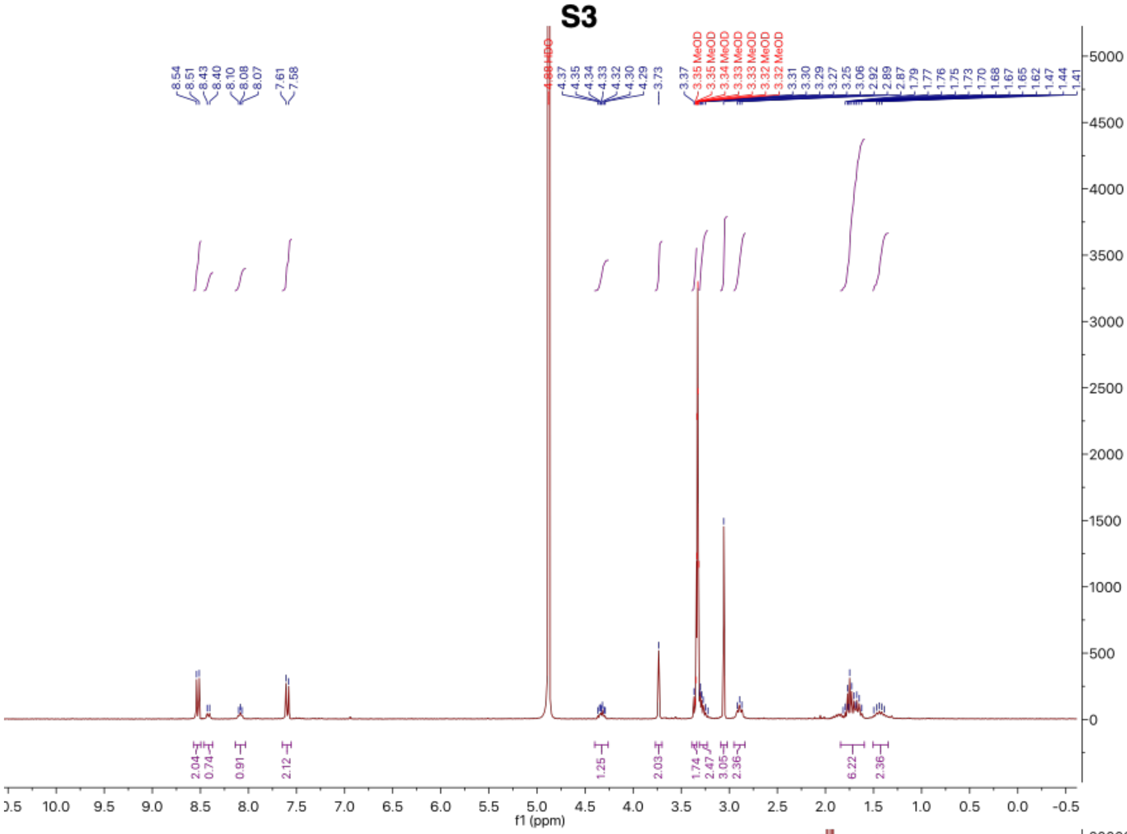
8.54
8.51
8.43
8.40
8.10
8.08
8.07
7.61
7.58
4.88 HDO
4.37
4.35
4.34
4.33
4.32
4.30
4.29
3.73
3.37
3.35 MeOD
3.35 MeOD
3.34 MeOD
3.33 MeOD
3.33 MeOD
3.32 MeOD
3.32 MeOD
3.31
3.30
3.29
3.27
3.25
3.06
2.92
2.89
2.87
1.79
1.77
1.76
1.75
1.73
1.70
1.68
1.67
1.65
1.62
1.47
1.44
1.41
2.04
0.74
0.91
2.12
1.25
2.03
1.74
2.47
3.05
2.36
6.22
2.36
f1 (ppm)

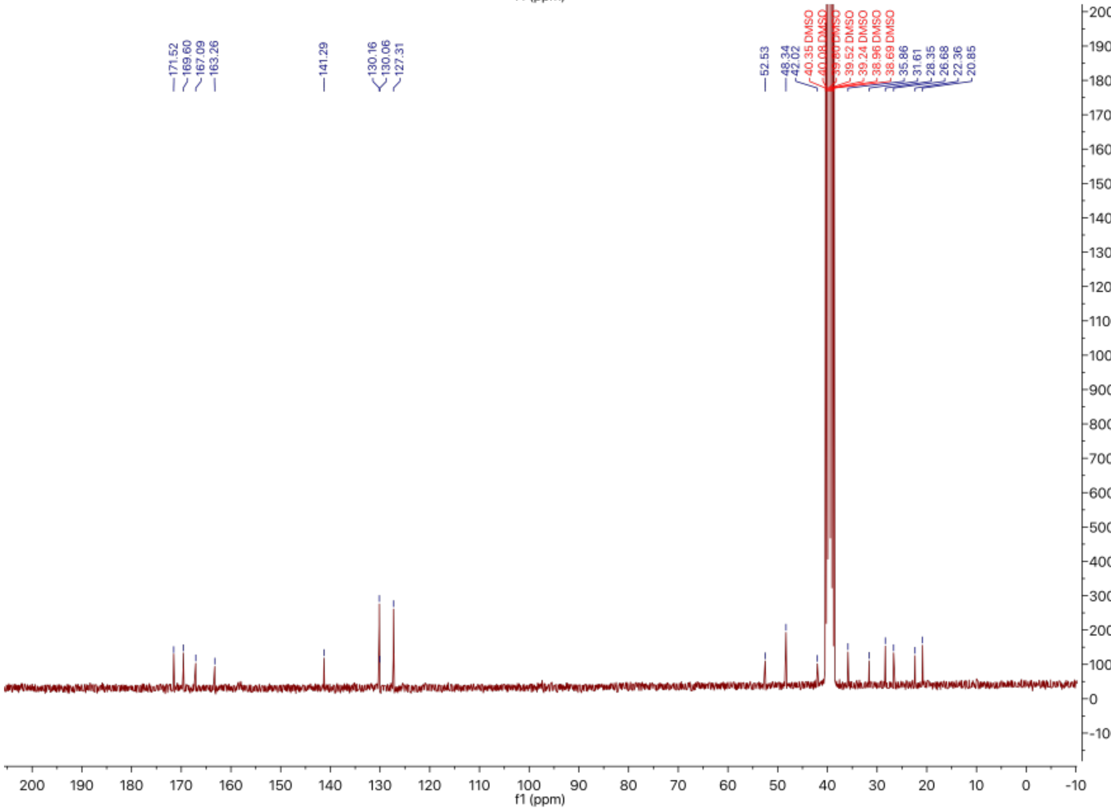
171.52
169.60
167.09
163.26
141.29
130.16
130.06
127.31
52.53
48.34
42.02
40.35 DMSO
40.08 DMSO
39.80 DMSO
39.52 DMSO
39.24 DMSO
38.96 DMSO
38.69 DMSO
35.86
31.61
28.35
26.68
22.36
20.85
f1 (ppm)

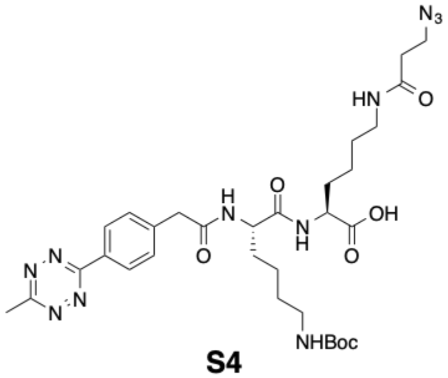

**S4**

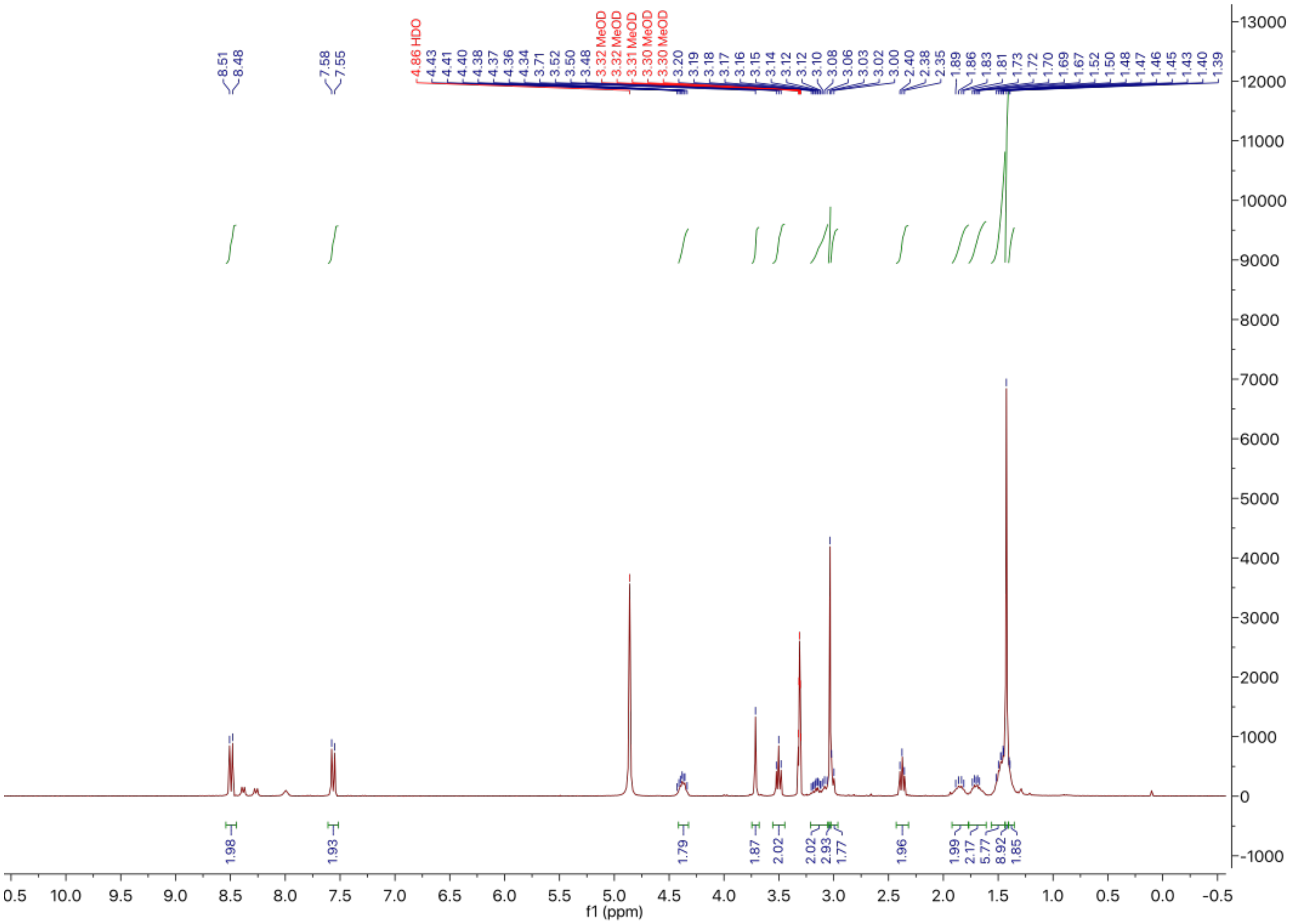

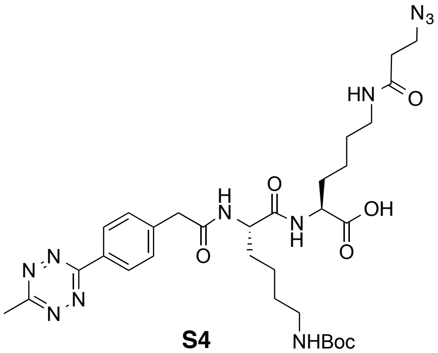

**S4**

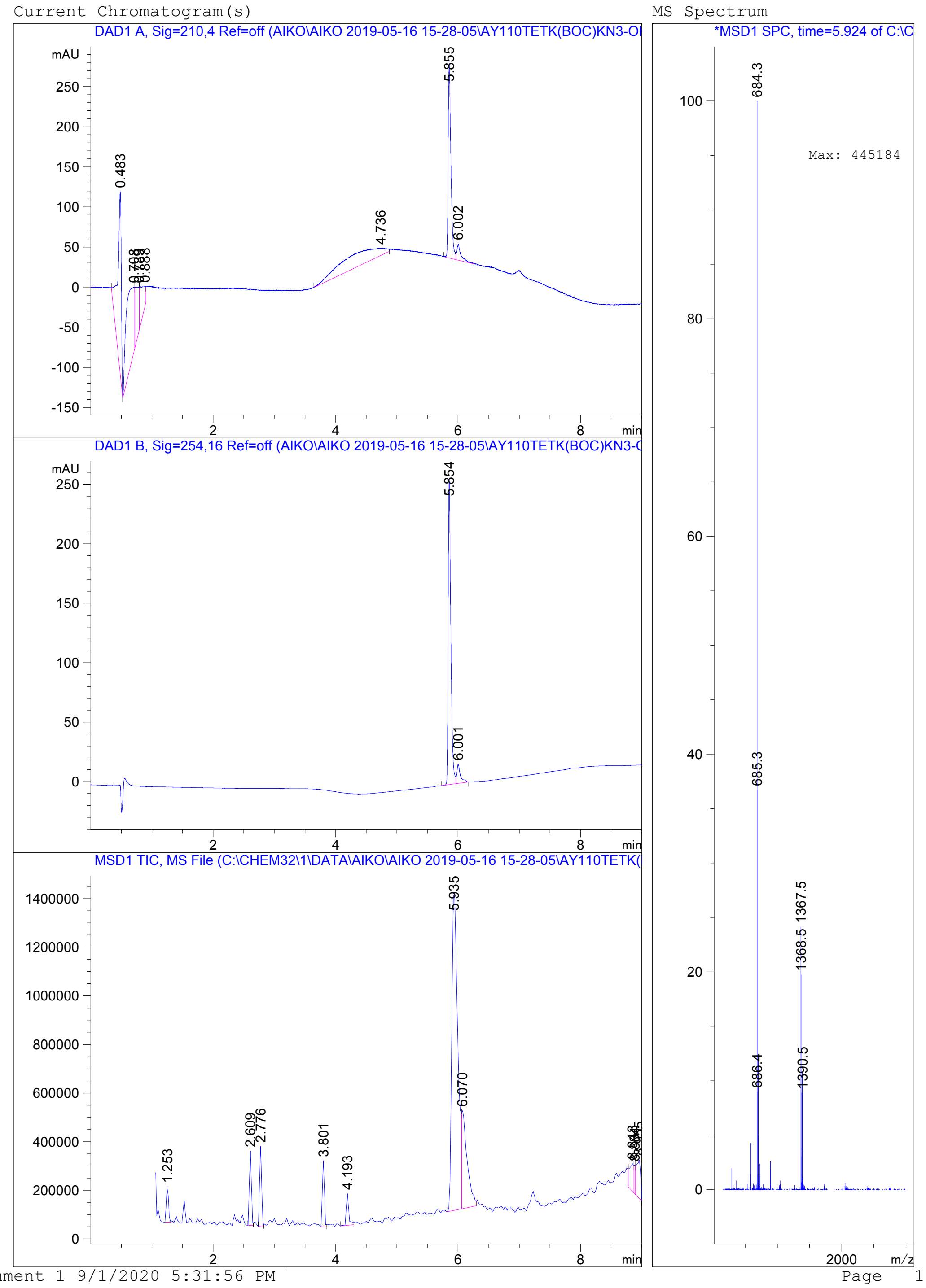

Instrument 1 9/1/2020 5:31:56 PM

Page 1 of 1

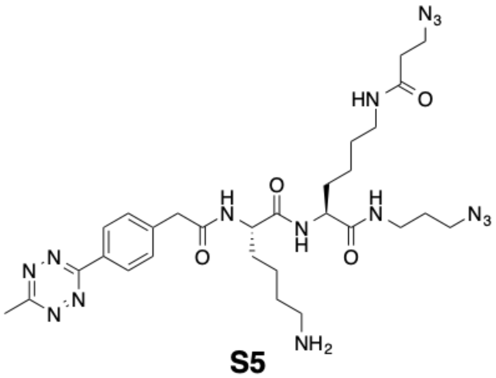
N3
HN
O
H
N
O
N
H
O
H
N
N3
N
N
N
N
O
NH2

**S5**

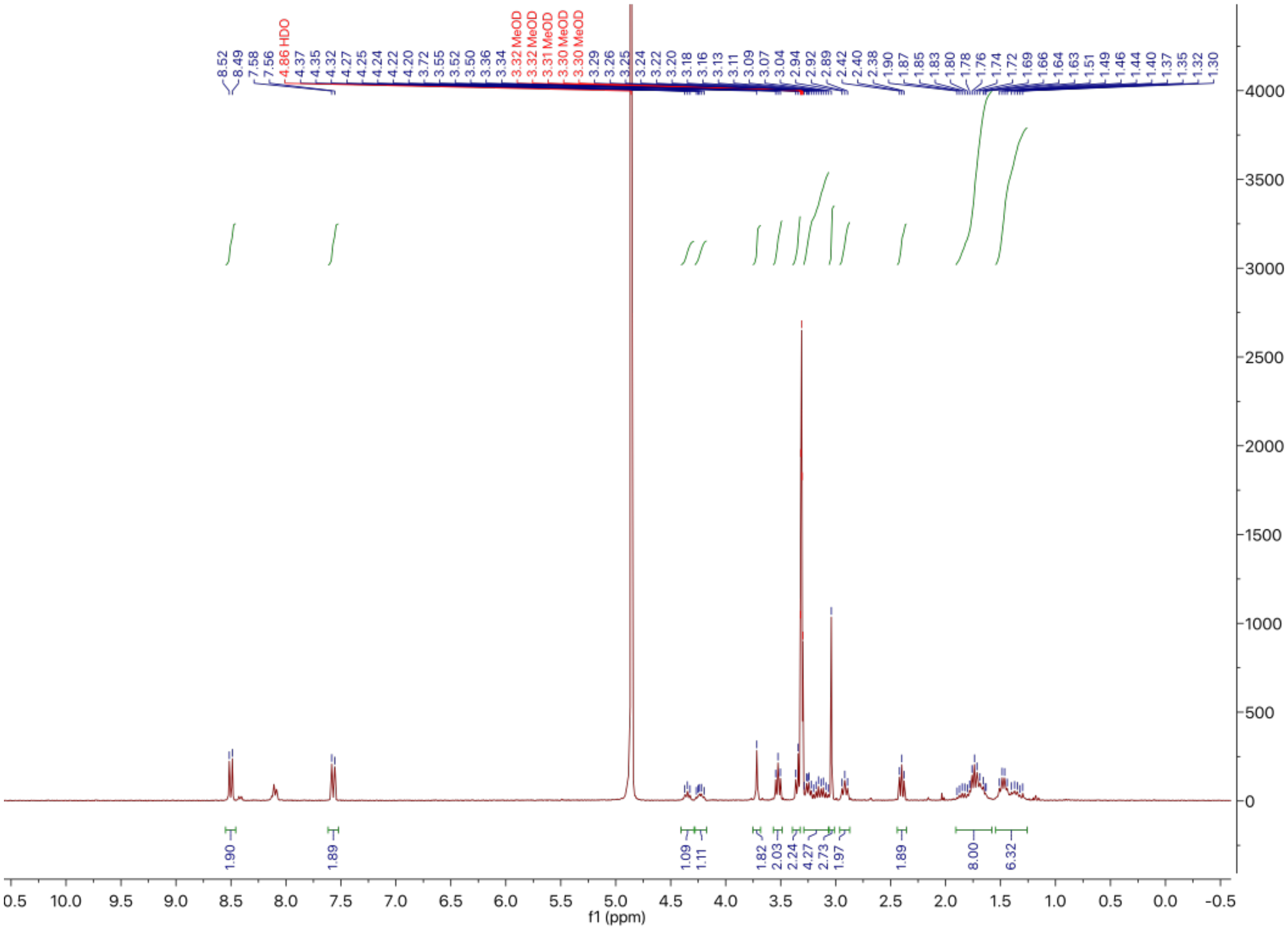
8.52
8.49
7.58
7.56
4.86 HDO
4.37
4.35
4.32
4.27
4.25
4.24
4.22
4.20
3.72
3.55
3.52
3.50
3.36
3.34
3.32 MeOD
3.32 MeOD
3.31 MeOD
3.30 MeOD
3.30 MeOD
3.29
3.26
3.25
3.24
3.22
3.20
3.18
3.16
3.13
3.11
3.09
3.07
3.04
2.94
2.92
2.89
2.42
2.40
2.38
1.90
1.87
1.85
1.83
1.80
1.78
1.76
1.74
1.72
1.69
1.66
1.64
1.63
1.51
1.49
1.46
1.44
1.40
1.37
1.35
1.32
1.30
1.90
1.89
1.09
1.11
1.82
2.03
2.24
4.27
2.73
1.97
1.89
8.00
6.32
0.5
10.0
9.5
9.0
8.5
8.0
7.5
7.0
6.5
6.0
5.5
5.0
4.5
4.0
3.5
3.0
2.5
2.0
1.5
1.0
0.5
0.0
-0.5
f1 (ppm)
4000
3500
3000
2500
2000
1500
1000
500
0

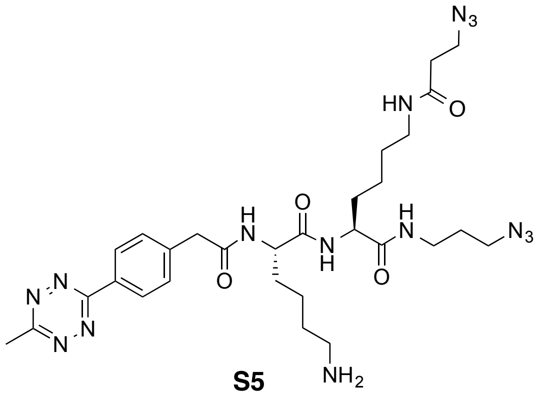
N3
HN
O
H
N
O
N
H
N
H
N3
N
N
N
N
O
NH2

**S5**

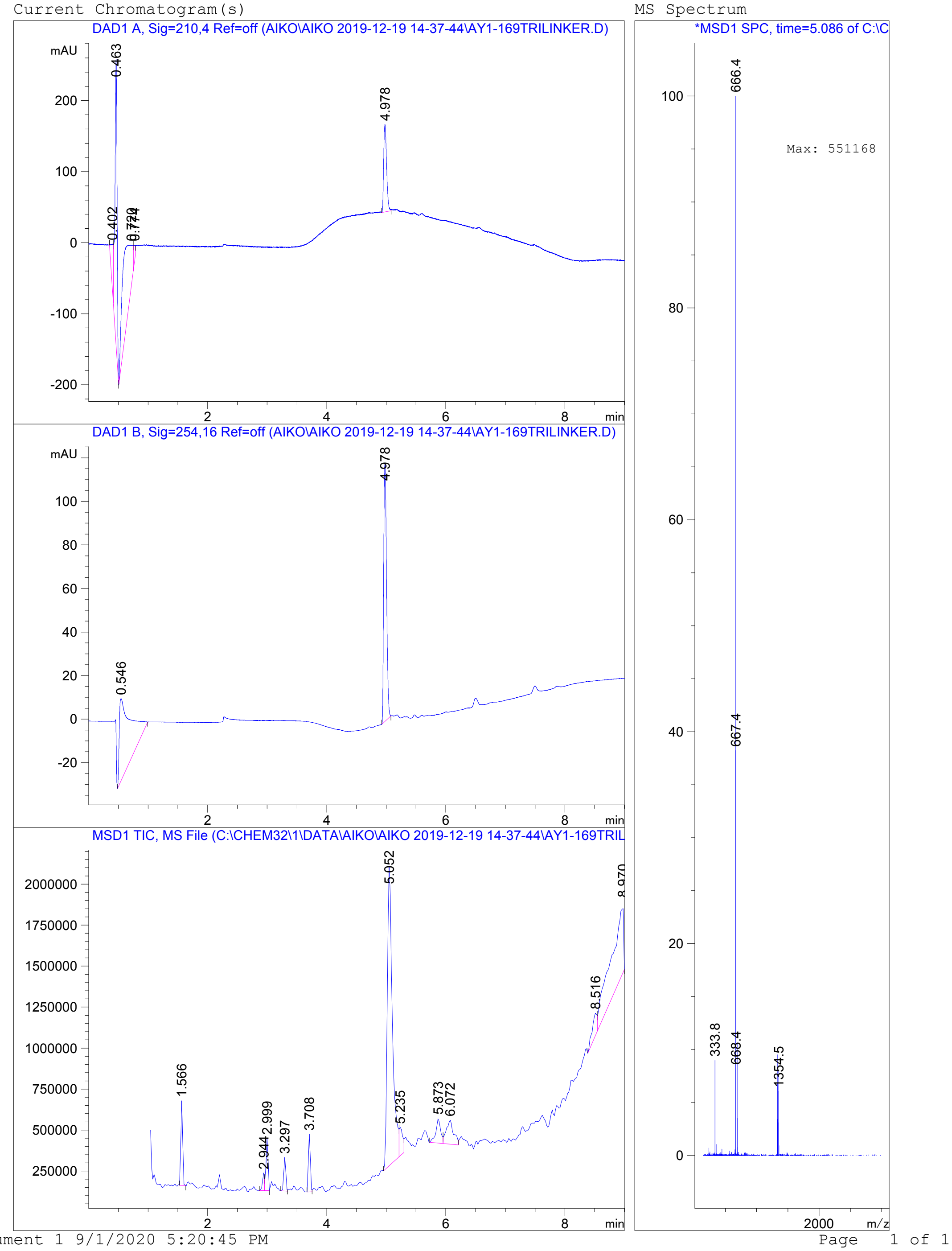
Current Chromatogram(s)
DAD1 A, Sig=210,4 Ref=off (AIKO\AIKO 2019-12-19 14-37-44\AY1-169TRILINKER.D)
mAU
0.463
4.978
0.402
min
DAD1 B, Sig=254,16 Ref=off (AIKO\AIKO 2019-12-19 14-37-44\AY1-169TRILINKER.D)
4.978
0.546
MSD1 TIC, MS File (C:\CHEM32\1\DATA\AIKO\AIKO 2019-12-19 14-37-44\AY1-169TRIL
5.052
1.566
2.944
2.999
3.297
3.708
5.235
5.873
6.072
8.516
MS Spectrum
*MSD1 SPC, time=5.086 of C:\C
666.4
Max: 551168
667.4
333.8
668.4
1354.5
m/z
Instrument 1 9/1/2020 5:20:45 PM
Page 1 of 1

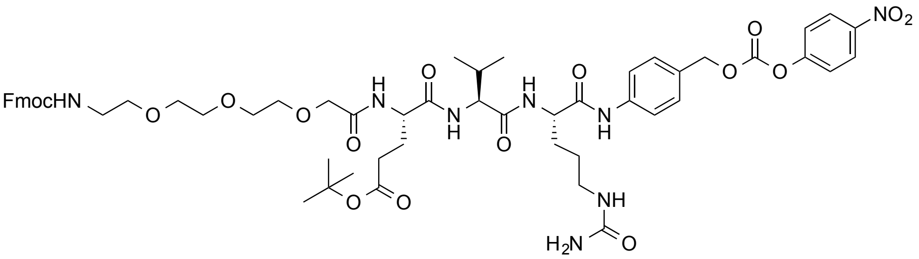
FmocHN
NO2
H2N
NH

**S6**

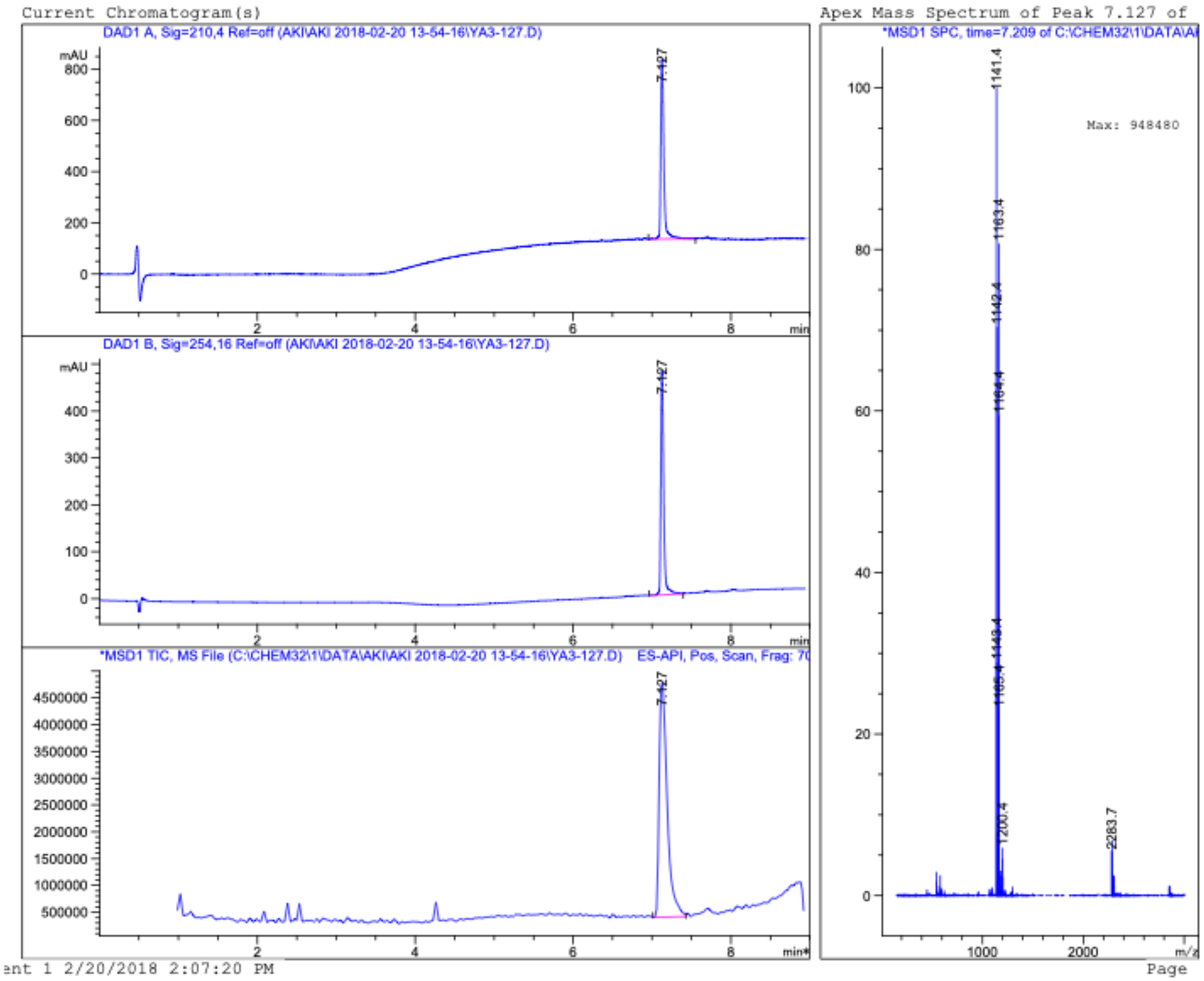
Current Chromatogram(s)
DAD1 A, Sig=210,4 Ref=off (AKI\AKI 2018-02-20 13-54-16\YA3-127.D)
mAU
7.127
DAD1 B, Sig=254,16 Ref=off (AKI\AKI 2018-02-20 13-54-16\YA3-127.D)
*MSD1 TIC, MS File (C:\CHEM32\1\DATA\AKI\AKI 2018-02-20 13-54-16\YA3-127.D) ES-API, Pos, Scan, Frag: 70
min
Apex Mass Spectrum of Peak 7.127 of
*MSD1 SPC, time=7.209 of C:\CHEM32\1\DATA\AI
Max: 948480
1141.4
1163.4
1142.4
1164.4
1143.4
1165.4
1200.4
2283.7
m/z
Page

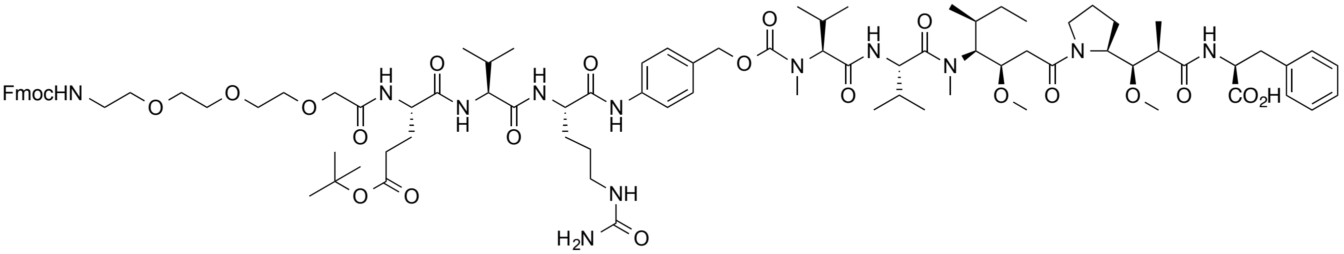
FmocHN
CO2H
H2N

**S7a**

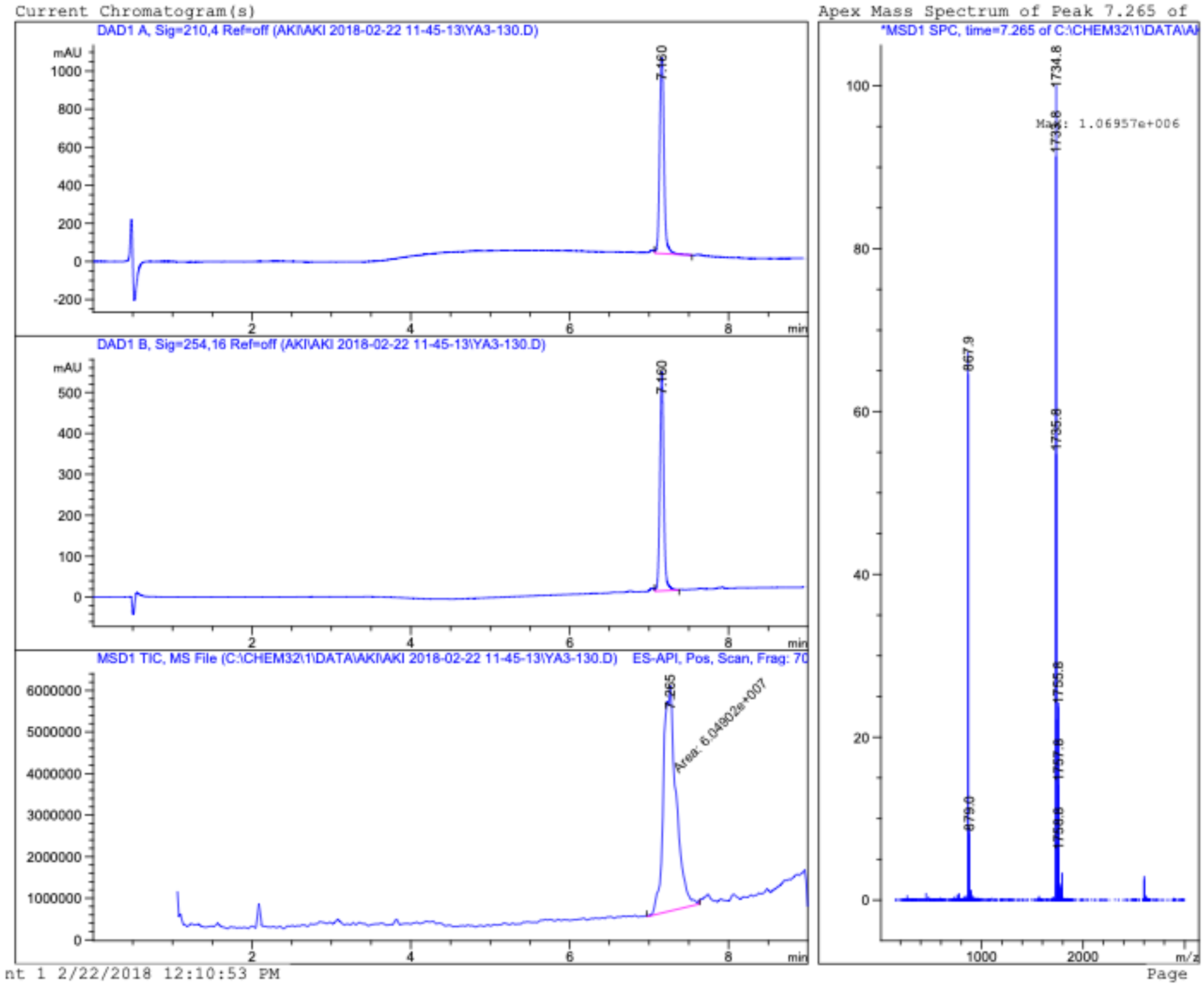
Current Chromatogram(s)
DAD1 A, Sig=210,4 Ref=off (AKI\AKI 2018-02-22 11-45-13\YA3-130.D)
mAU
7.160
DAD1 B, Sig=254,16 Ref=off (AKI\AKI 2018-02-22 11-45-13\YA3-130.D)
MSD1 TIC, MS File (C:\CHEM32\1\DATA\AKI\AKI 2018-02-22 11-45-13\YA3-130.D) ES-API, Pos, Scan, Frag: 70
7.265
Area: 6.04902e+007
min
Apex Mass Spectrum of Peak 7.265 of
*MSD1 SPC, time=7.265 of C:\CHEM32\1\DATA\AI
Max: 1.06957e+006
1734.8
1735.8
1755.8
867.9
879.0
m/z
nt 1 2/22/2018 12:10:53 PM
Page

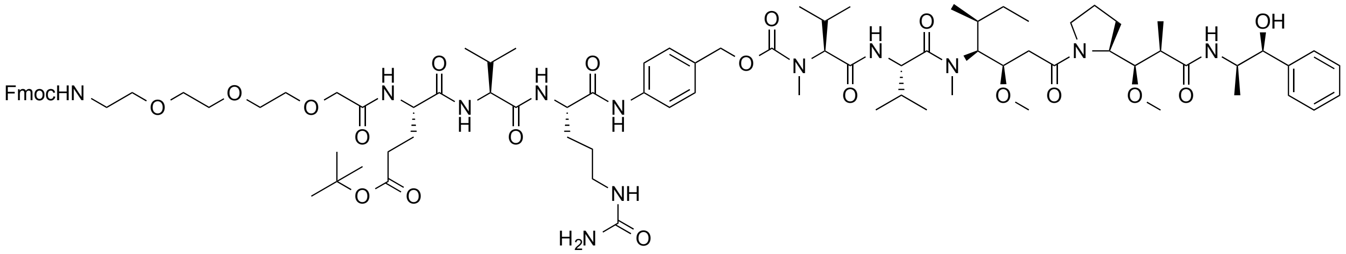
FmocHN
H2N
OH

**S7b**

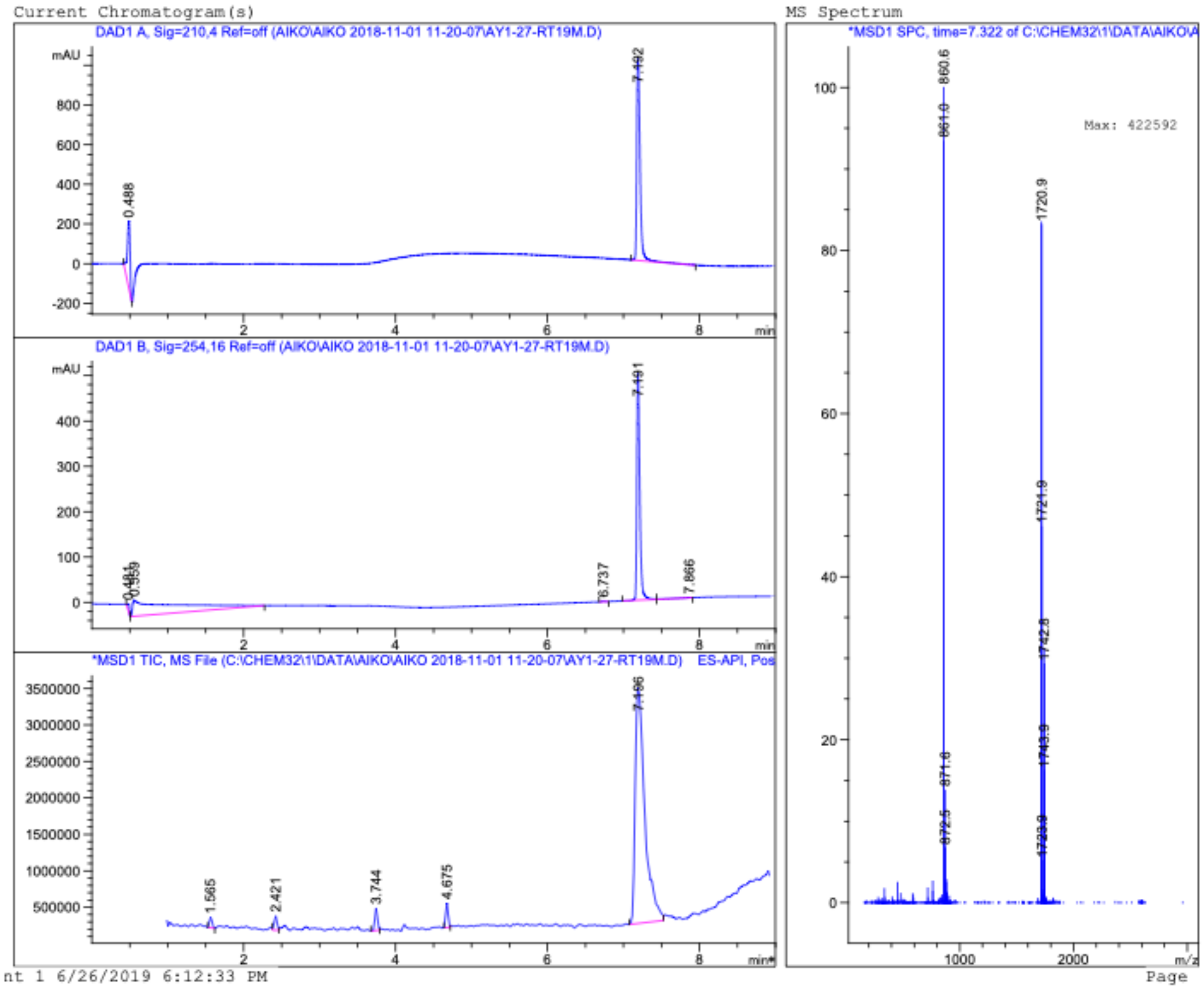
Current Chromatogram(s)
DAD1 A, Sig=210,4 Ref=off (AIKO\AIKO 2018-11-01 11-20-07\AY1-27-RT19M.D)
mAU
0.488
7.192
min
DAD1 B, Sig=254,16 Ref=off (AIKO\AIKO 2018-11-01 11-20-07\AY1-27-RT19M.D)
6.737
7.191
7.866
*MSD1 TIC, MS File (C:\CHEM32\1\DATA\AIKO\AIKO 2018-11-01 11-20-07\AY1-27-RT19M.D) ES-API, Pos
1.565
2.421
3.744
4.675
MS Spectrum
*MSD1 SPC, time=7.322 of C:\CHEM32\1\DATA\AIKO\A
Max: 422592
860.6
861.0
871.8
1720.9
1721.9
1742.8
m/z
nt 1 6/26/2019 6:12:33 PM
Page

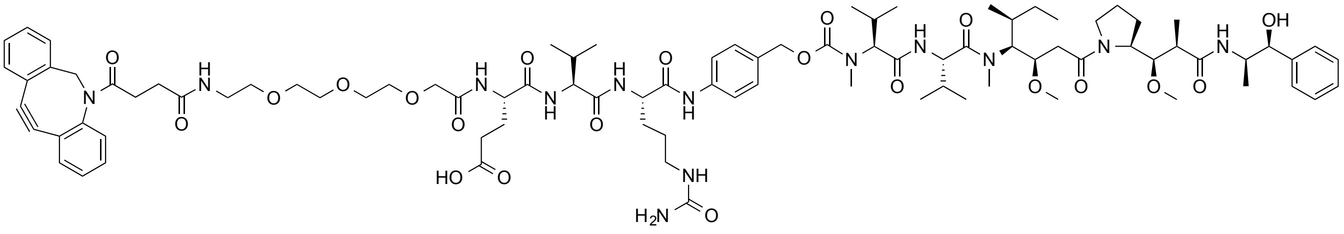

**S8b**

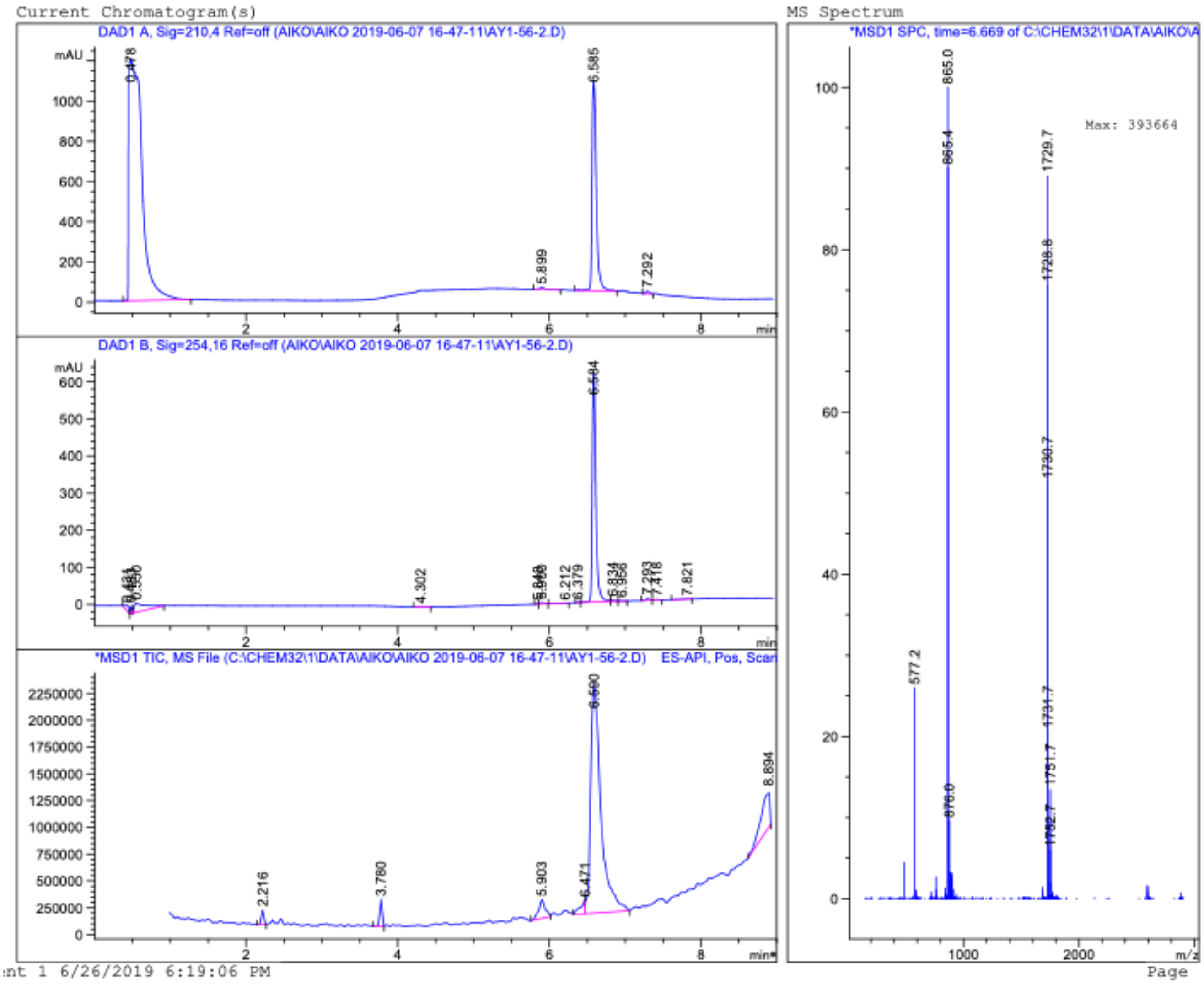
Current Chromatogram(s)
DAD1 A, Sig=210,4 Ref=off (AIKO\AIKO 2019-06-07 16-47-11\AY1-56-2.D)
DAD1 B, Sig=254,16 Ref=off (AIKO\AIKO 2019-06-07 16-47-11\AY1-56-2.D)
*MSD1 TIC, MS File (C:\CHEM32\1\DATA\AIKO\AIKO 2019-06-07 16-47-11\AY1-56-2.D) ES-API, Pos, Scan
MS Spectrum
*MSD1 SPC, time=6.669 of C:\CHEM32\1\DATA\AIKO\A
Max: 393664
865.0
1729.7
577.2
6.585
6.584
6.590
ent 1 6/26/2019 6:19:06 PM
Page

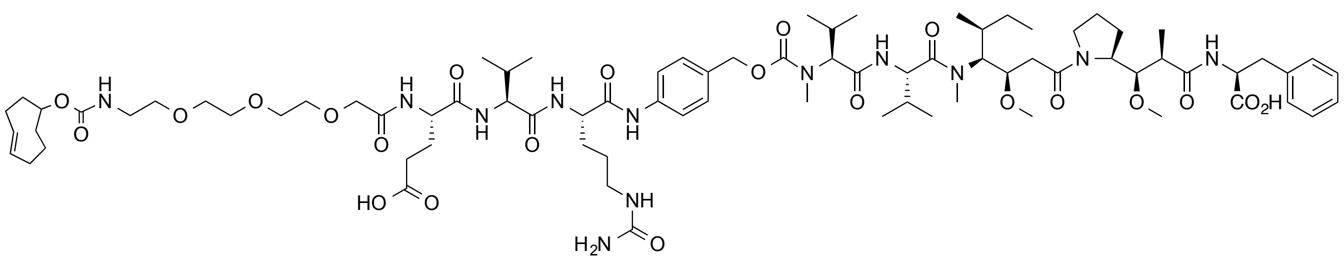

**S9a**

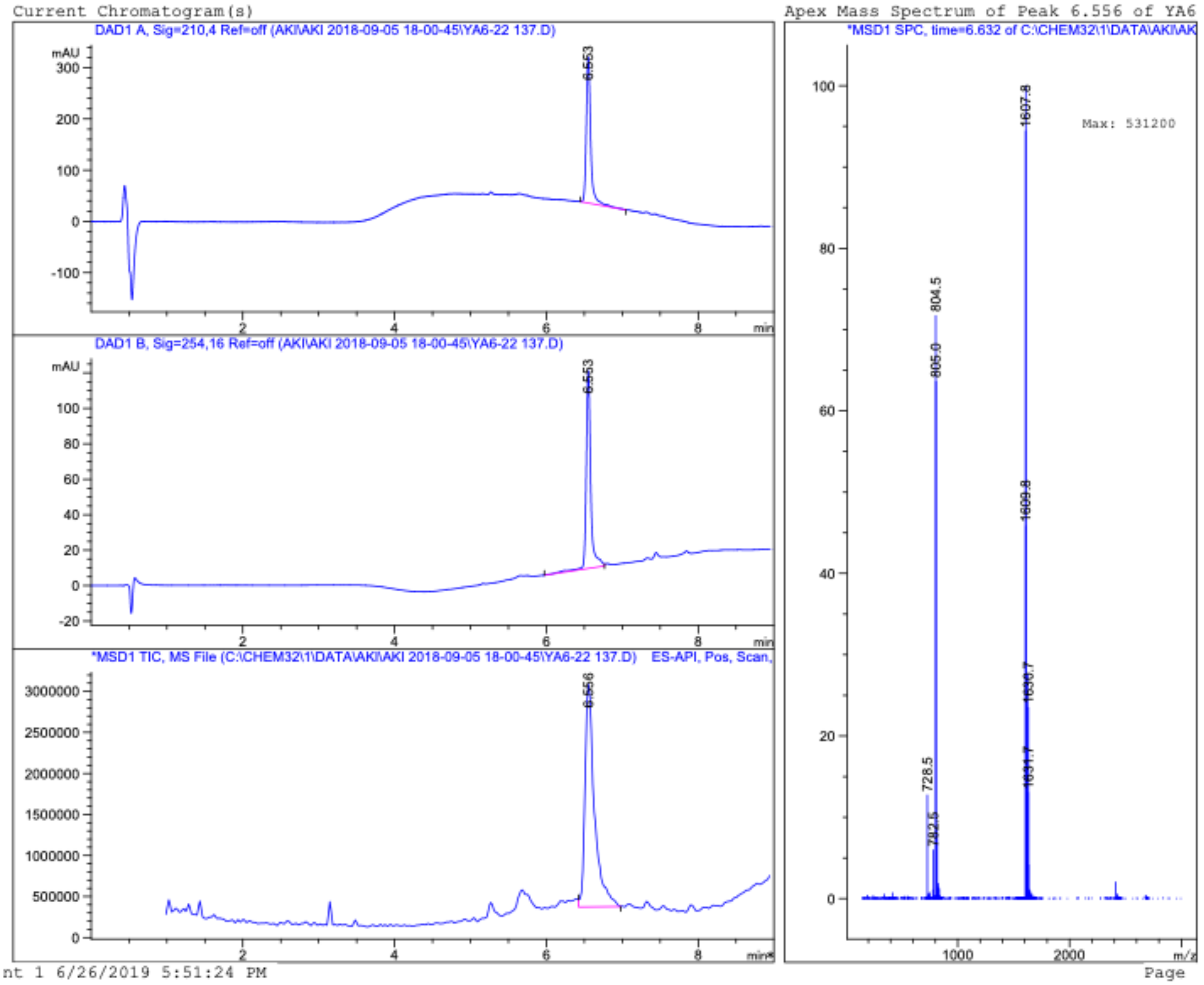

Current Chromatogram(s)
DAD1 A, Sig=210,4 Ref=off (AKI\AKI 2018-09-05 18-00-45\YA6-22 137.D)
mAU
6.553
DAD1 B, Sig=254,16 Ref=off (AKI\AKI 2018-09-05 18-00-45\YA6-22 137.D)
*MSD1 TIC, MS File (C:\CHEM32\1\DATA\AKI\AKI 2018-09-05 18-00-45\YA6-22 137.D) ES-API, Pos, Scan,
6.556
min
nt 1 6/26/2019 5:51:24 PM
Apex Mass Spectrum of Peak 6.556 of YA6
*MSD1 SPC, time=6.632 of C:\CHEM32\1\DATA\AKI\AK
Max: 531200
1607.8
804.5
805.0
1609.8
1630.7
1631.7
728.5
782.6
m/z
Page

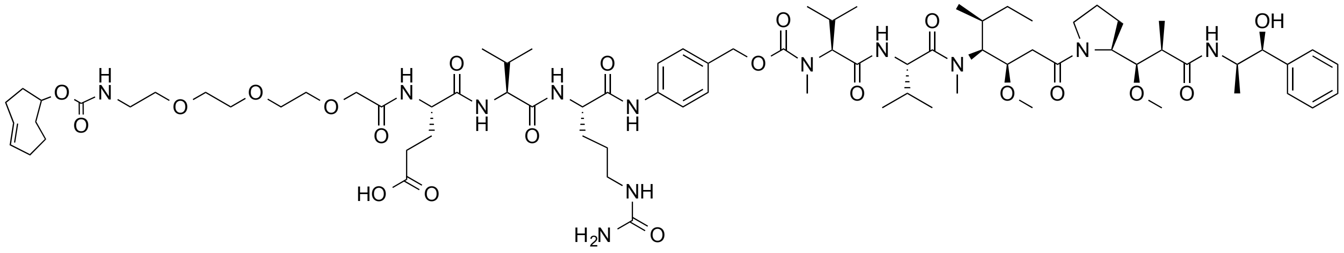

**S9b**

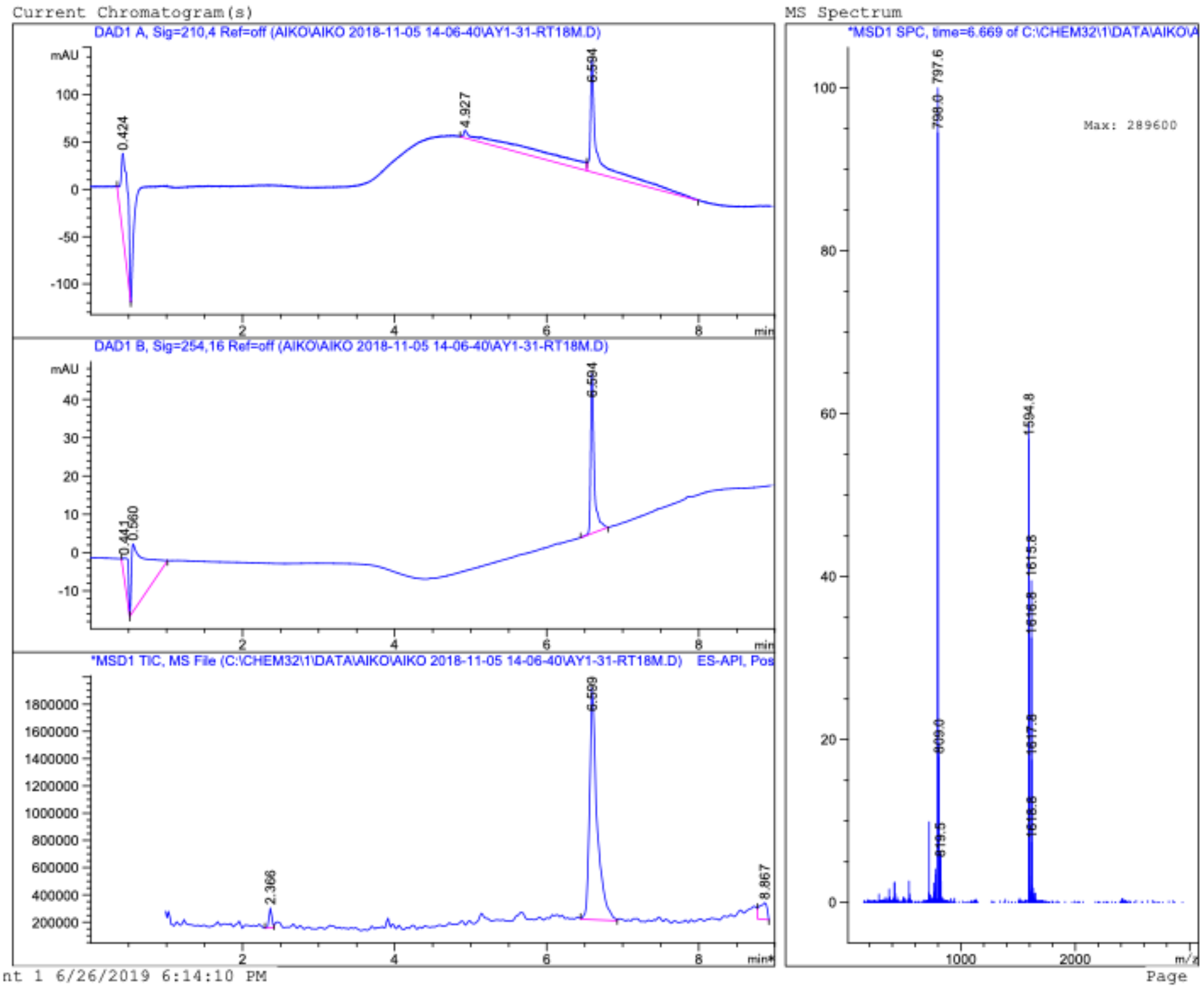
Current Chromatogram(s)
DAD1 A, Sig=210,4 Ref=off (AIKO\AIKO 2018-11-05 14-06-40\AY1-31-RT18M.D)
mAU
0.424
4.927
6.594
min
DAD1 B, Sig=254,16 Ref=off (AIKO\AIKO 2018-11-05 14-06-40\AY1-31-RT18M.D)
0.441
0.560
6.594
*MSD1 TIC, MS File (C:\CHEM32\1\DATA\AIKO\AIKO 2018-11-05 14-06-40\AY1-31-RT18M.D) ES-API, Pos
2.366
6.599
8.867
MS Spectrum
*MSD1 SPC, time=6.669 of C:\CHEM32\1\DATA\AIKO\A
Max: 289600
797.6
798.0
809.0
819.5
1594.8
1615.8
1616.8
1617.8
1618.8
m/z
nt 1 6/26/2019 6:14:10 PM
Page

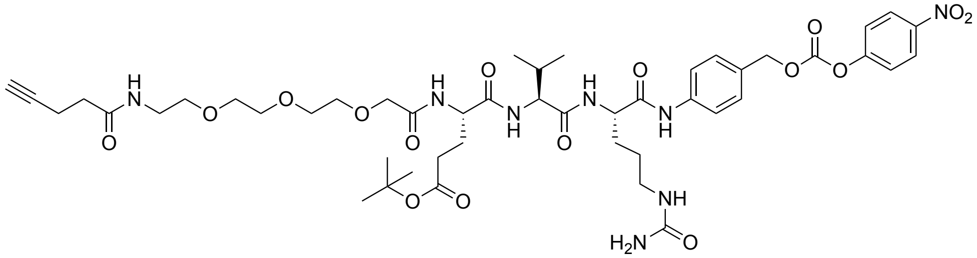
NO2
H
N
O
O
O
H
N
O
N
H
H
N
O
N
H
O
O
O
NH
H2N
O

**S10a**

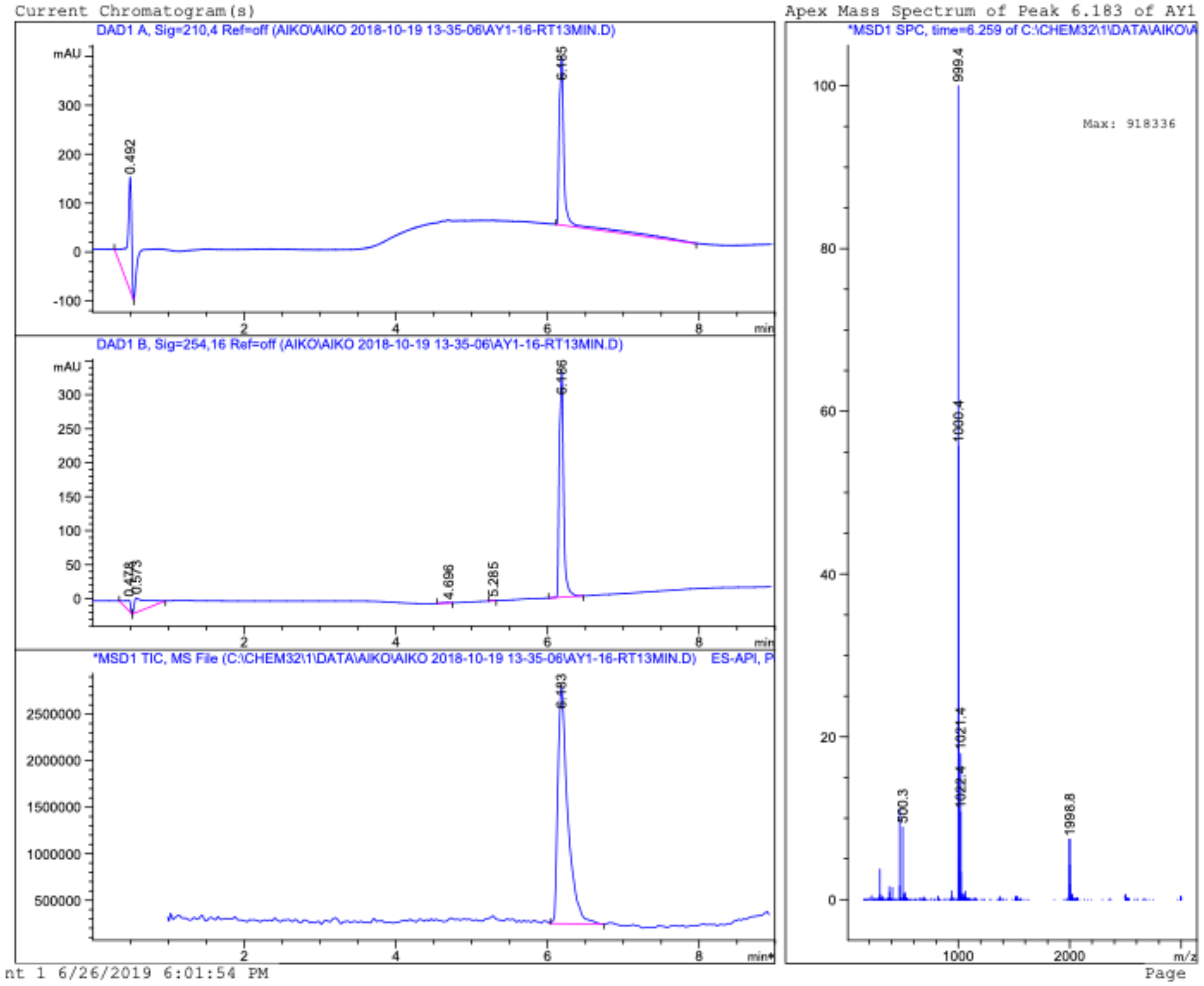
Current Chromatogram(s)
DAD1 A, Sig=210,4 Ref=off (AIKO\AIKO 2018-10-19 13-35-06\AY1-16-RT13MIN.D)
0.492
6.185
DAD1 B, Sig=254,16 Ref=off (AIKO\AIKO 2018-10-19 13-35-06\AY1-16-RT13MIN.D)
0.478
0.573
4.696
5.285
6.186
*MSD1 TIC, MS File (C:\CHEM32\1\DATA\AIKO\AIKO 2018-10-19 13-35-06\AY1-16-RT13MIN.D) ES-API, P
6.183
Apex Mass Spectrum of Peak 6.183 of AY1
*MSD1 SPC, time=6.259 of C:\CHEM32\1\DATA\AIKO\A
Max: 918336
999.4
1000.4
1021.4
1022.4
500.3
1998.8
nt 1 6/26/2019 6:01:54 PM
Page

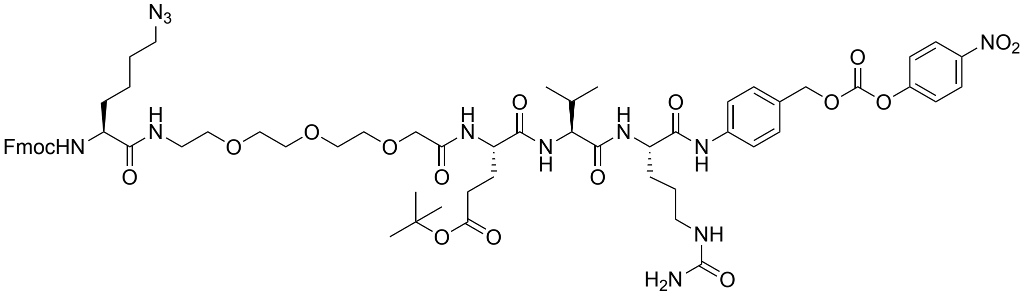
N3
NO2
FmocHN
H2N

**S10b**

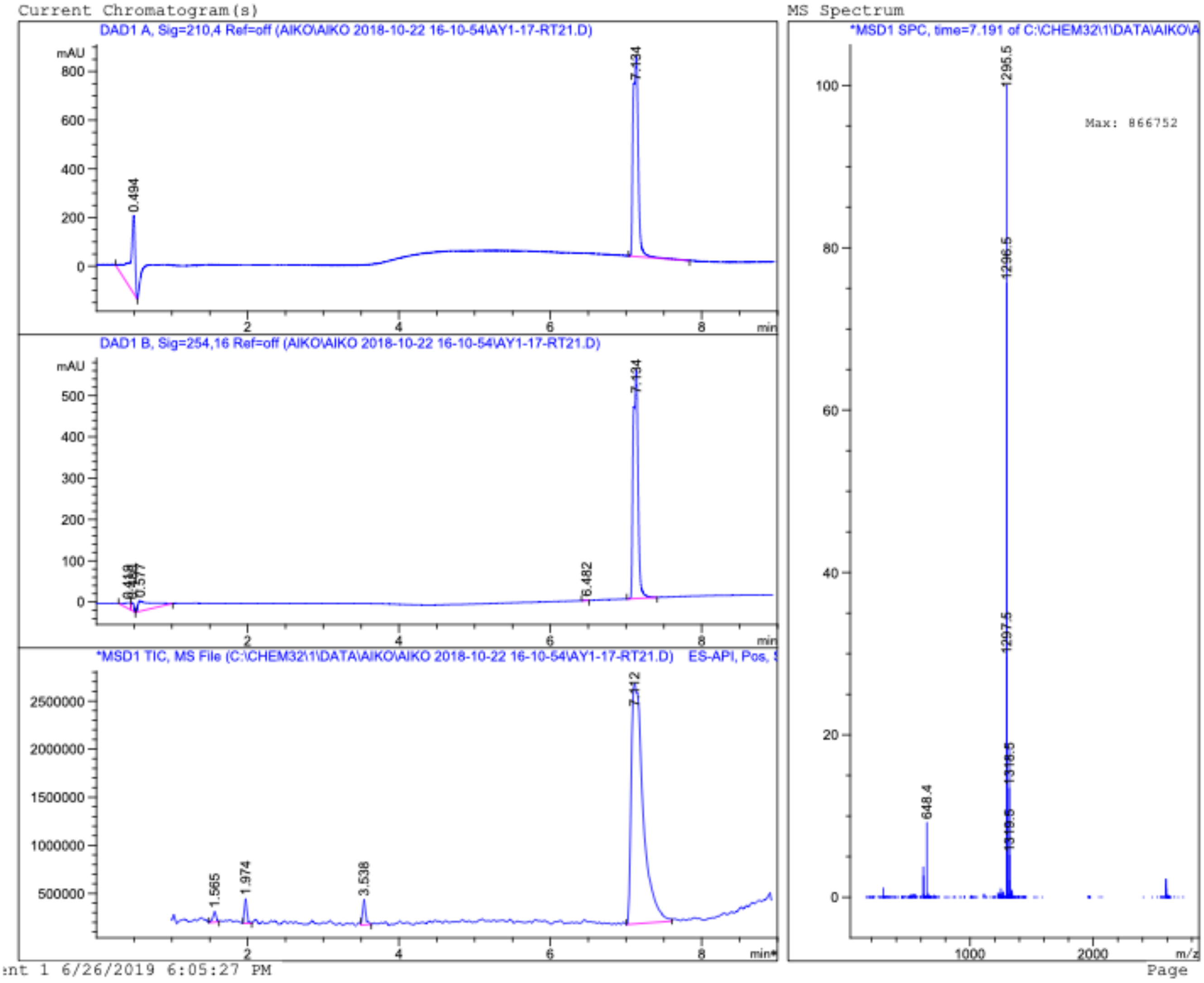
Current Chromatogram(s)
DAD1 A, Sig=210,4 Ref=off (AIKO\AIKO 2018-10-22 16-10-54\AY1-17-RT21.D)
mAU
7.134
0.494
DAD1 B, Sig=254,16 Ref=off (AIKO\AIKO 2018-10-22 16-10-54\AY1-17-RT21.D)
7.134
0.577
6.482
*MSD1 TIC, MS File (C:\CHEM32\1\DATA\AIKO\AIKO 2018-10-22 16-10-54\AY1-17-RT21.D) ES-API, Pos,
7.112
1.565
1.974
3.538
min
MS Spectrum
*MSD1 SPC, time=7.191 of C:\CHEM32\1\DATA\AIKO\A
Max: 866752
1295.5
1296.5
1297.5
1318.5
648.4
m/z
nt 1 6/26/2019 6:05:27 PM
Page

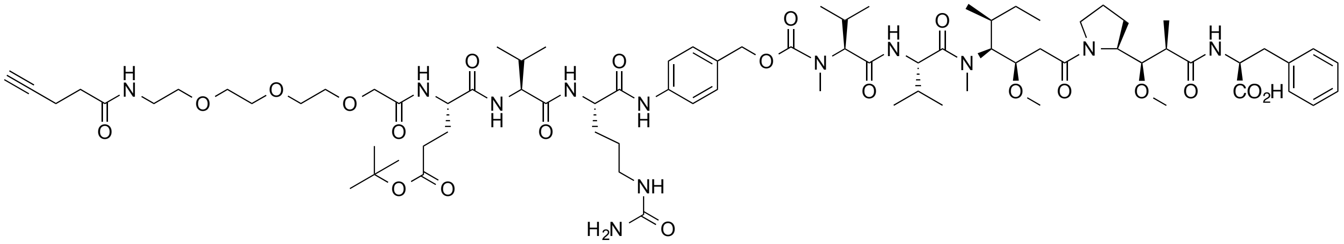

**S11a**

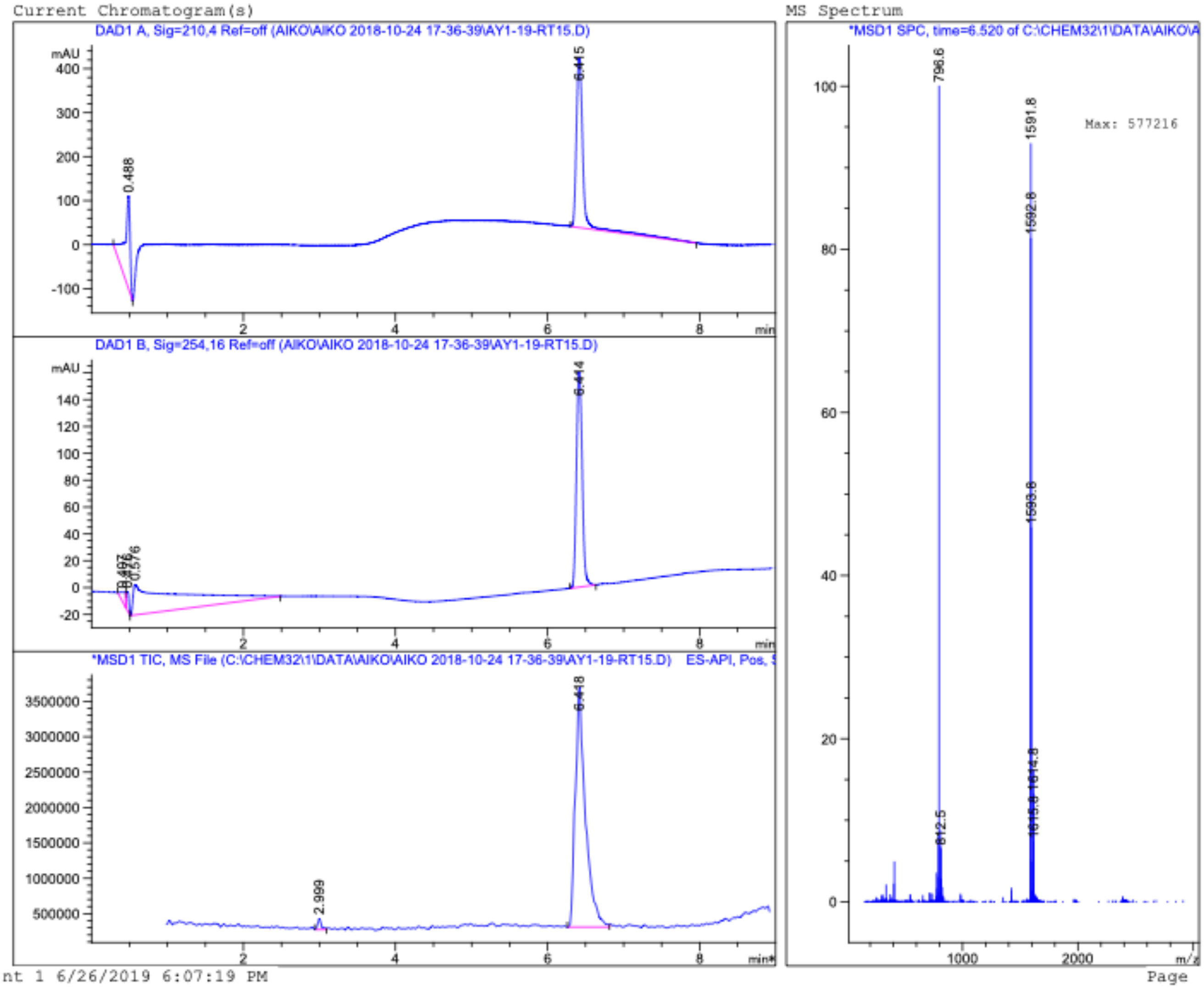
Current Chromatogram(s)
DAD1 A, Sig=210,4 Ref=off (AIKO\AIKO 2018-10-24 17-36-39\AY1-19-RT15.D)
mAU
6.415
0.488
DAD1 B, Sig=254,16 Ref=off (AIKO\AIKO 2018-10-24 17-36-39\AY1-19-RT15.D)
mAU
6.414
0.576
*MSD1 TIC, MS File (C:\CHEM32\1\DATA\AIKO\AIKO 2018-10-24 17-36-39\AY1-19-RT15.D) ES-API, Pos,
6.448
2.999
min
MS Spectrum
*MSD1 SPC, time=6.520 of C:\CHEM32\1\DATA\AIKO\A
796.6
1591.8
1592.8
1593.8
1614.8
812.5
Max: 577216
m/z
nt 1 6/26/2019 6:07:19 PM
Page

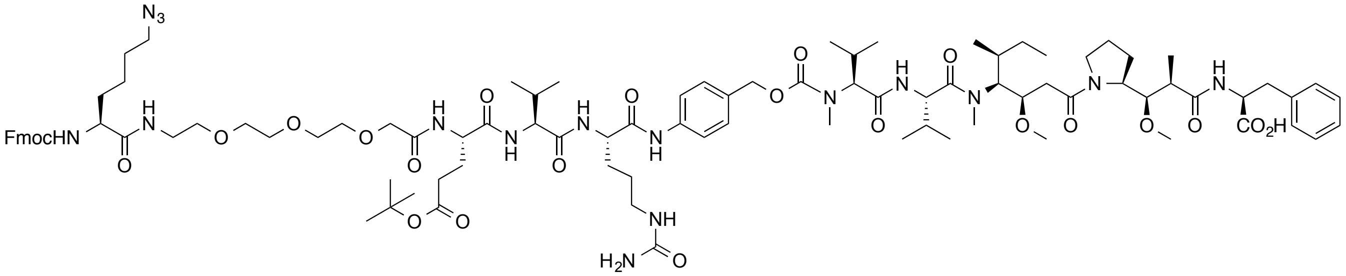
N3
FmocHN
H
N
O
O
O
O
O
H
N
N
H
O
O
H
N
N
H
O
NH
H2N
O
O
N
H
O
N
N
O
O
N
N
O
O
N
H
CO2H

**S11b**

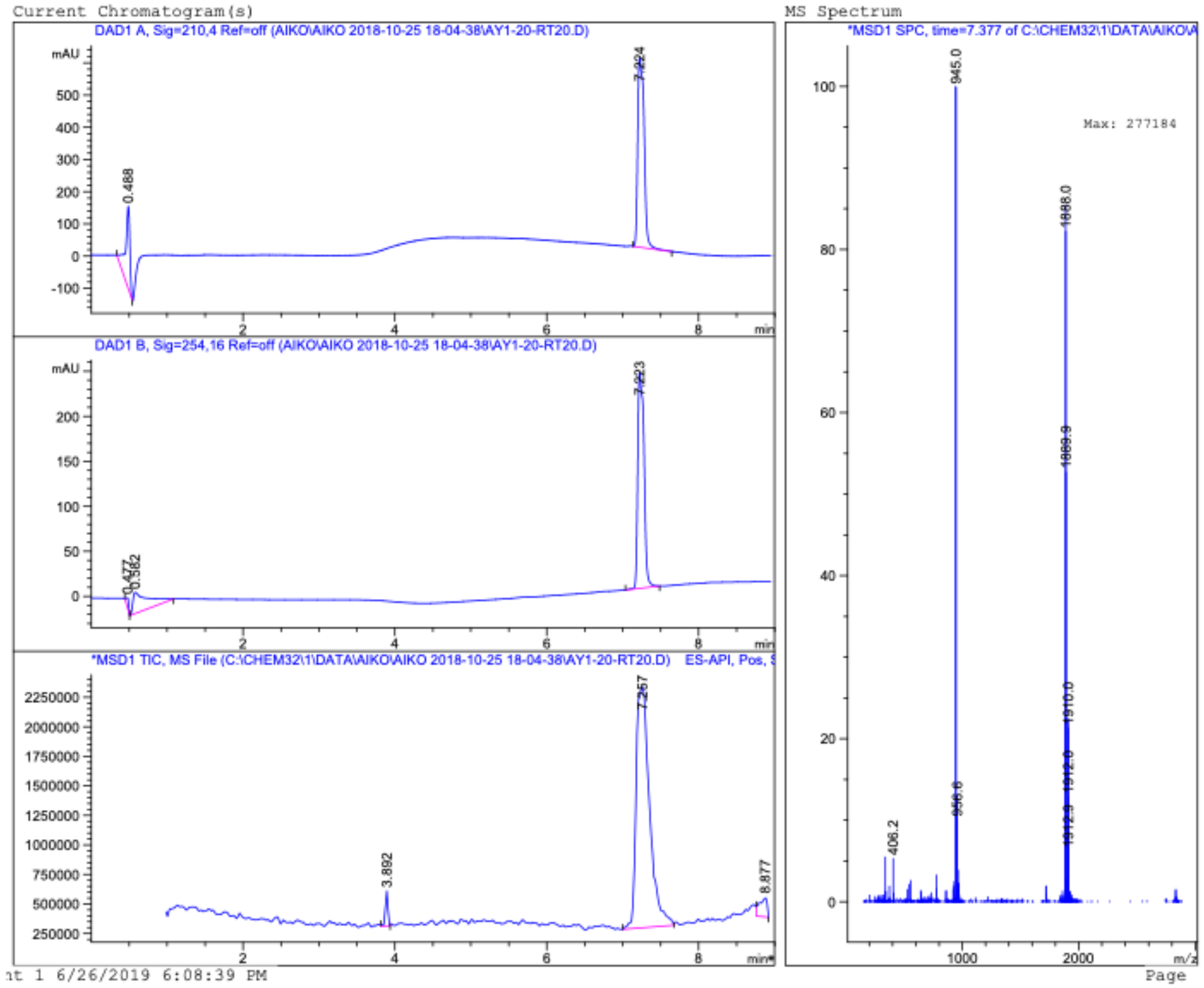
Current Chromatogram(s)
DAD1 A, Sig=210,4 Ref=off (AIKO\AIKO 2018-10-25 18-04-38\AY1-20-RT20.D)
mAU
0.488
7.224
DAD1 B, Sig=254,16 Ref=off (AIKO\AIKO 2018-10-25 18-04-38\AY1-20-RT20.D)
7.223
*MSD1 TIC, MS File (C:\CHEM32\1\DATA\AIKO\AIKO 2018-10-25 18-04-38\AY1-20-RT20.D) ES-API, Pos,
3.892
7.257
8.877
min
MS Spectrum
*MSD1 SPC, time=7.377 of C:\CHEM32\1\DATA\AIKO\
945.0
Max: 277184
1888.0
406.2
956.6
1910.0
m/z
6/26/2019 6:08:39 PM
Page

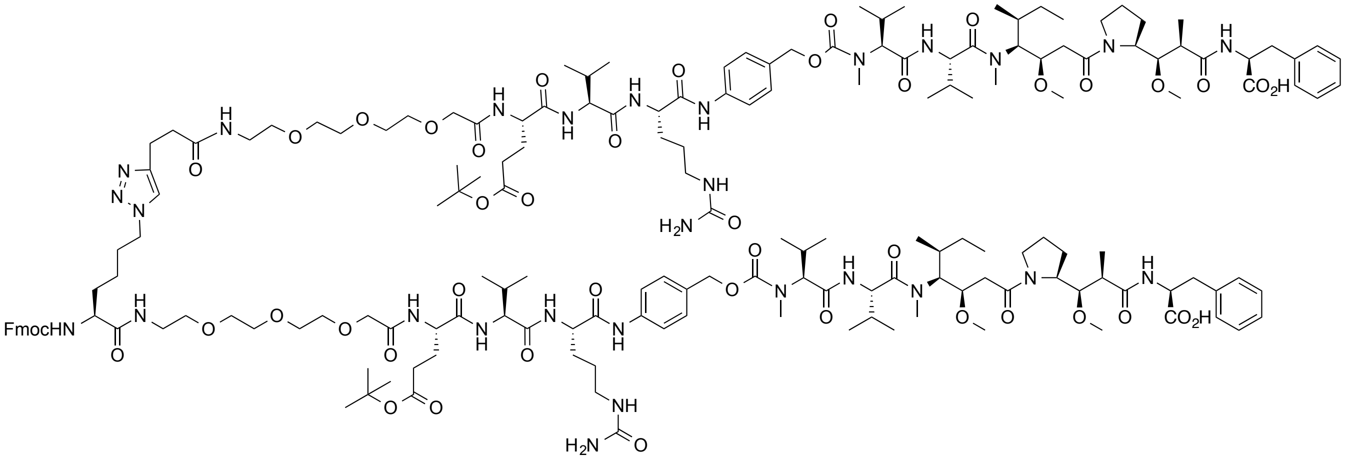
FmocHN
$H_2N$
$CO_2H$

**S12**

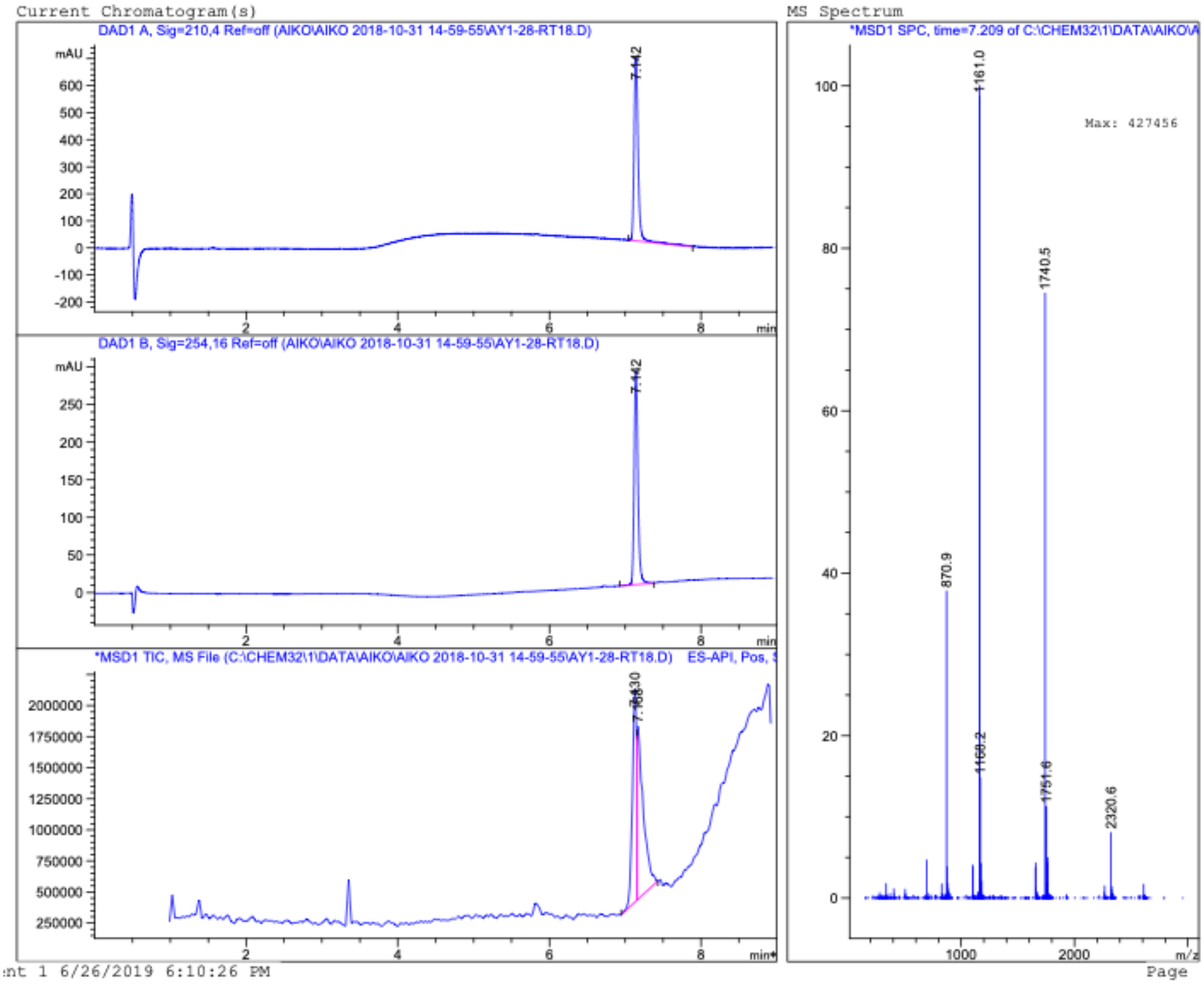
Current Chromatogram(s)
DAD1 A, Sig=210,4 Ref=off (AIKO\AIKO 2018-10-31 14-59-55\AY1-28-RT18.D)
7.142
DAD1 B, Sig=254,16 Ref=off (AIKO\AIKO 2018-10-31 14-59-55\AY1-28-RT18.D)
7.142
*MSD1 TIC, MS File (C:\CHEM32\1\DATA\AIKO\AIKO 2018-10-31 14-59-55\AY1-28-RT18.D) ES-API, Pos,
MS Spectrum
*MSD1 SPC, time=7.209 of C:\CHEM32\1\DATA\AIKO\A
Max: 427456
1161.0
1740.5
870.9
1168.2
1751.6
2320.6
m/z
min
mAU
nt 1 6/26/2019 6:10:26 PM
Page

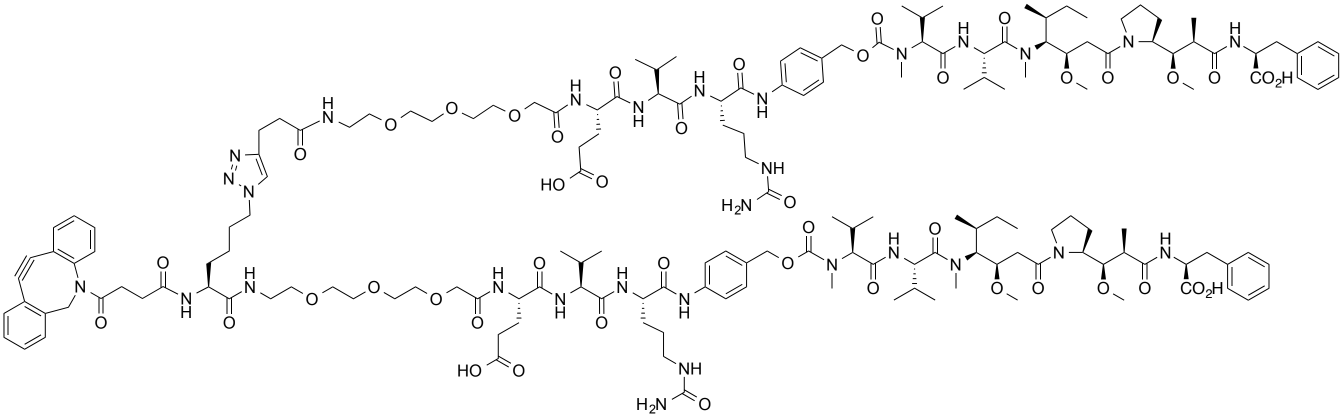

**S13**

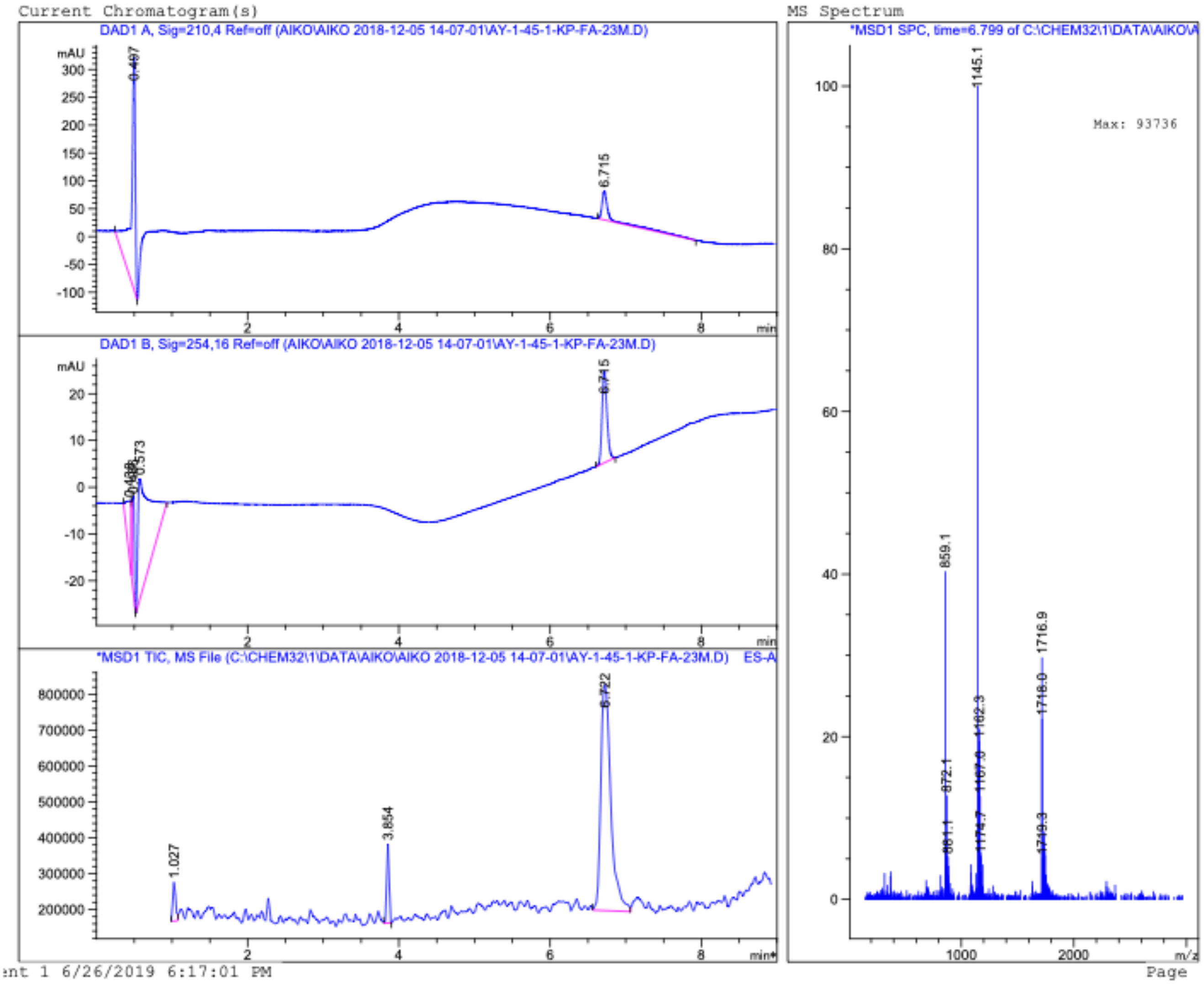
Current Chromatogram(s)
DAD1 A, Sig=210,4 Ref=off (AIKO\AIKO 2018-12-05 14-07-01\AY-1-45-1-KP-FA-23M.D)
6.715
DAD1 B, Sig=254,16 Ref=off (AIKO\AIKO 2018-12-05 14-07-01\AY-1-45-1-KP-FA-23M.D)
6.745
0.573
*MSD1 TIC, MS File (C:\CHEM32\1\DATA\AIKO\AIKO 2018-12-05 14-07-01\AY-1-45-1-KP-FA-23M.D) ES-A
1.027
3.854
6.722
MS Spectrum
*MSD1 SPC, time=6.799 of C:\CHEM32\1\DATA\AIKO\A
Max: 93736
1145.1
859.1
872.1
1162.3
1716.9
1718.0
1749.3
nt 1 6/26/2019 6:17:01 PM
Page

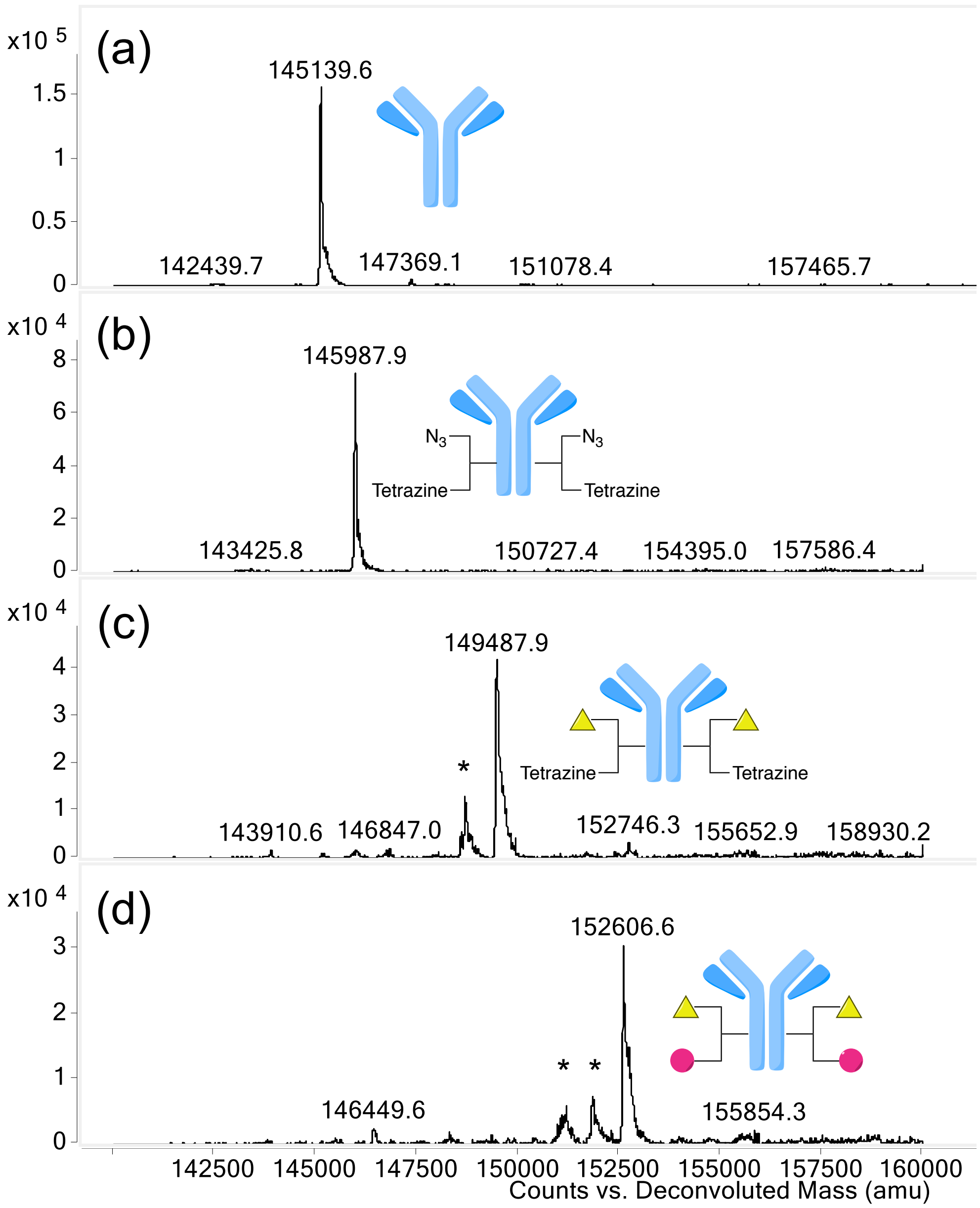

Deconvoluted ESI-mass spectra of light chain of aglycosylated N297A anti-HER2 mAb (a), mAb–azido-methyltetrazine di-arm linker conjugate (b), linker conjugate after DBCO–azide click reaction (c), and MMAE/F 2+2 ADCs (d). Asterisk (*) indicates fragment ion(s) detected in ESI-MS analysis.

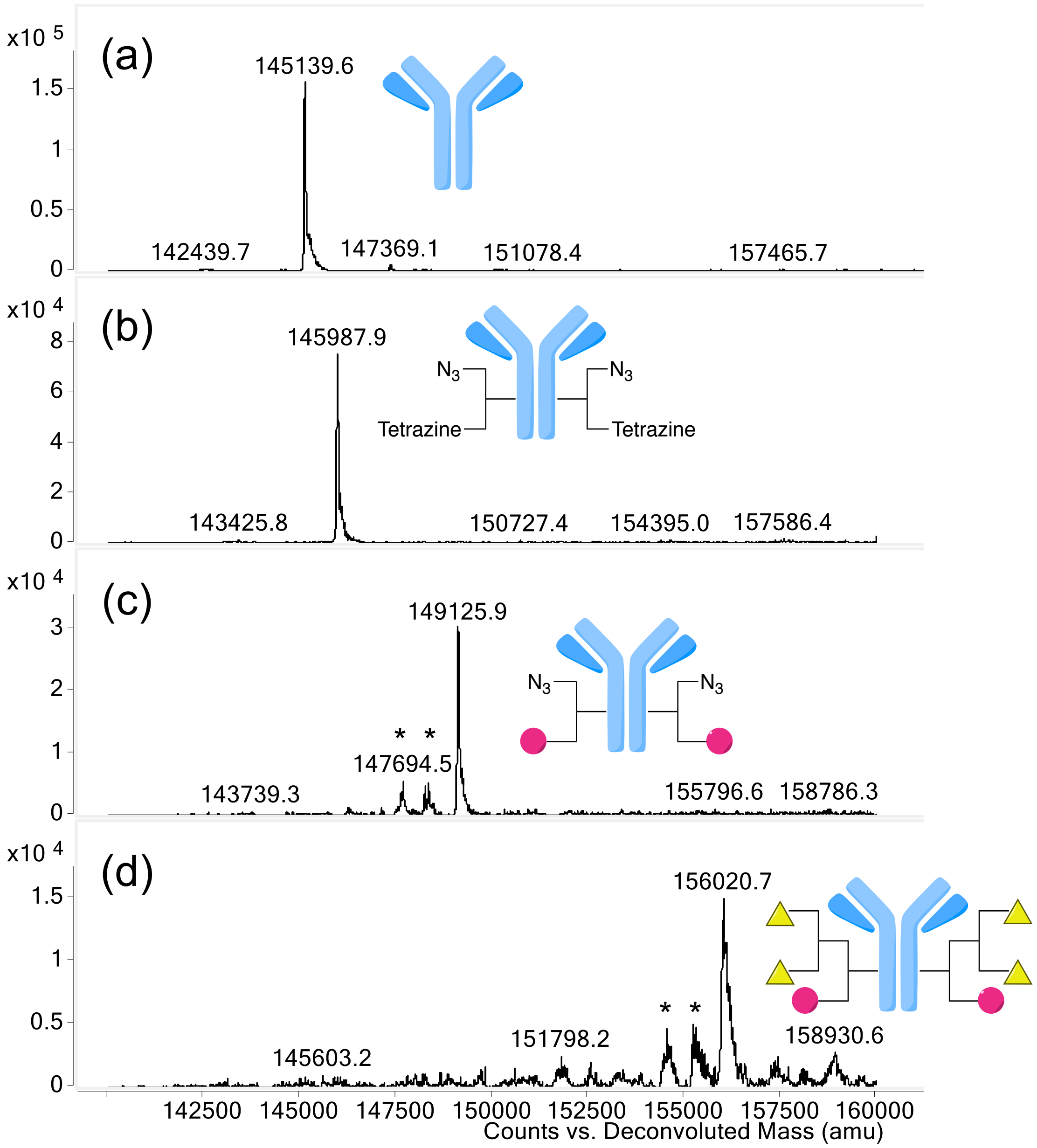

Deconvoluted ESI-mass spectra of light chain of aglycosylated N297A anti-HER2 mAb (a), mAb–azido-methyltetrazine di-arm linker conjugate (b), linker conjugate after TCO–methyltetrazine click reaction (c), and MMAE/F 2+4 ADC (d). Asterisk (*) indicates fragment ion(s) detected in ESI-MS analysis.

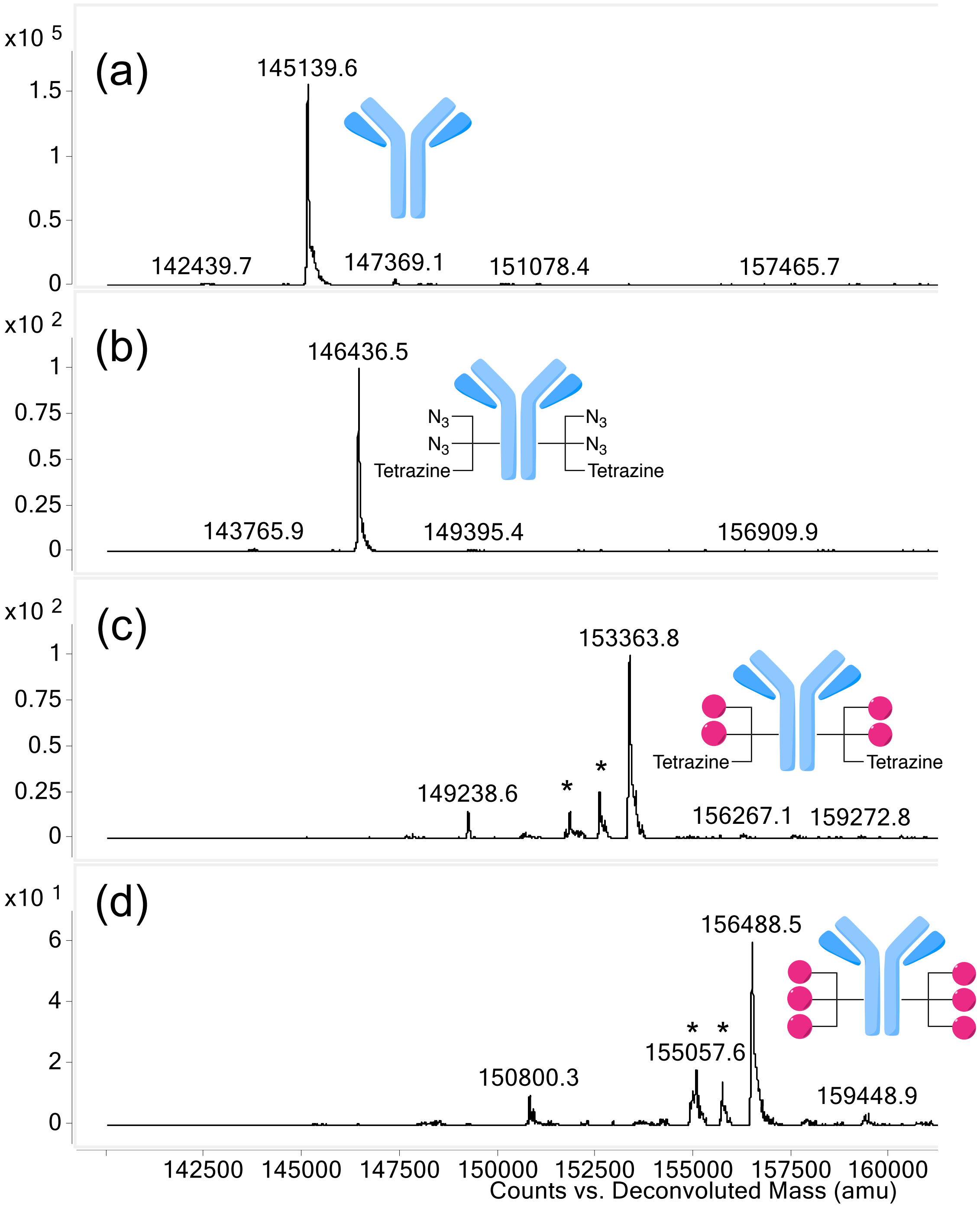

Deconvoluted ESI-mass spectra of light chain of aglycosylated N297A anti-HER2 mAb (a), mAb–diazido-methyltetrazine tri-arm linker conjugate (b), linker conjugate after DBCO–azide click reaction (c), and MMAE 6 ADC (d). Asterisk (*) indicates fragment ion(s) detected in ESI-MS analysis.

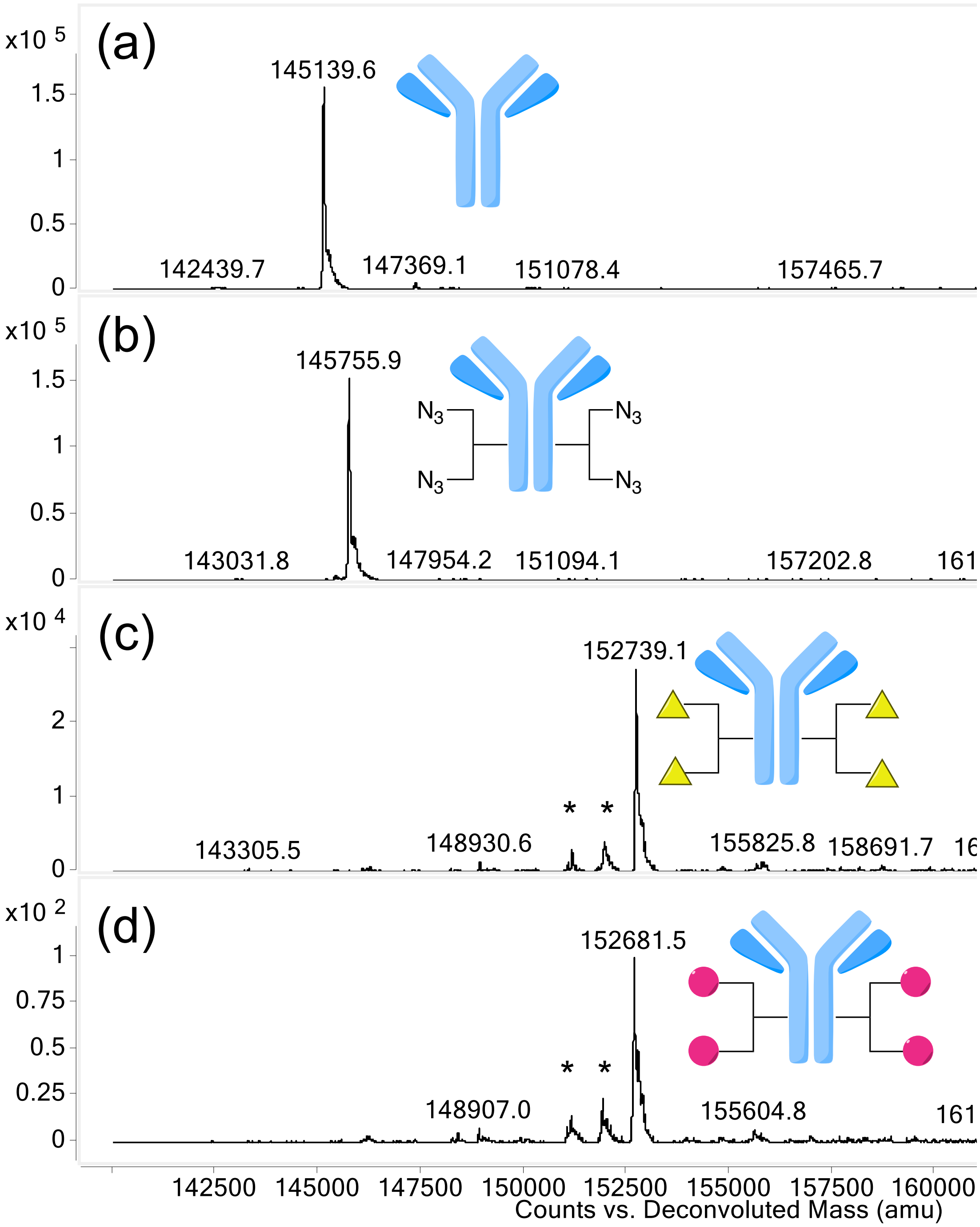

Deconvoluted ESI-mass spectra of light chain of aglycosylated N297A anti-HER2 mAb (a), mAb–diazide linker conjugate (b), MMAF DAR 4 ADC (c), and MMAE DAR 4 ADC (d). Asterisk (*) indicates fragment ion(s) detected in ESI-MS analysis.

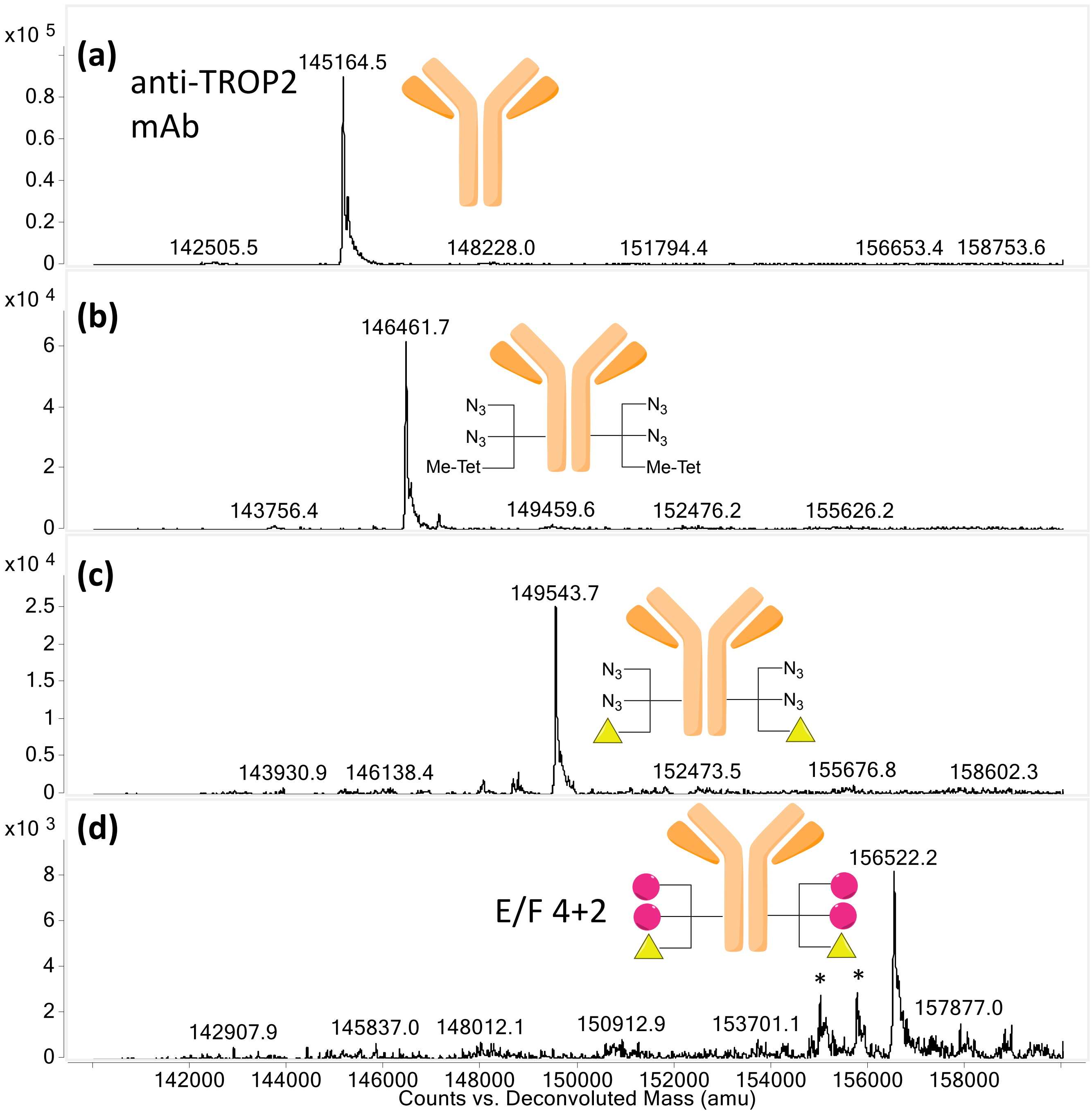

Deconvoluted ESI-mass spectra of light chain of aglycosylated N297A anti-TROP2 mAb (a), mAb–diazido-methyltetrazine tri-arm linker conjugate (b), linker conjugate after DBCO–azide click reaction (c), and MMAE/F 4+2 ADCs (d). Asterisk (*) indicates fragment ion(s) detected in ESI-MS analysis.

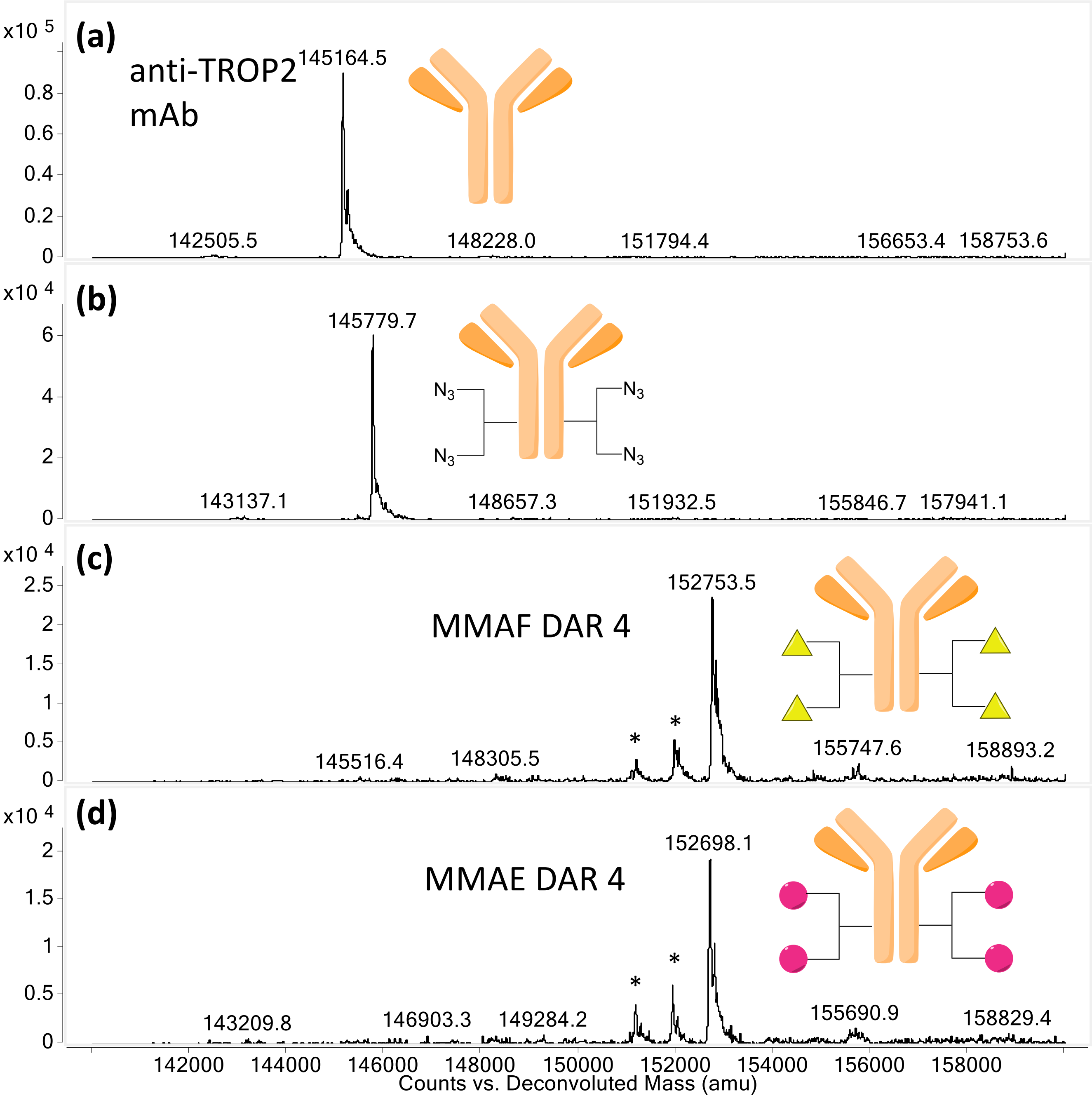

Deconvoluted ESI-mass spectra of light chain of aglycosylated N297A anti-TROP2 mAb (a), mAb–diazide linker conjugate (b), MMAF DAR 4 ADC (c), and MMAE DAR 4 ADC (d). Asterisk (*) indicates fragment ion(s) detected in ESI-MS analysis.

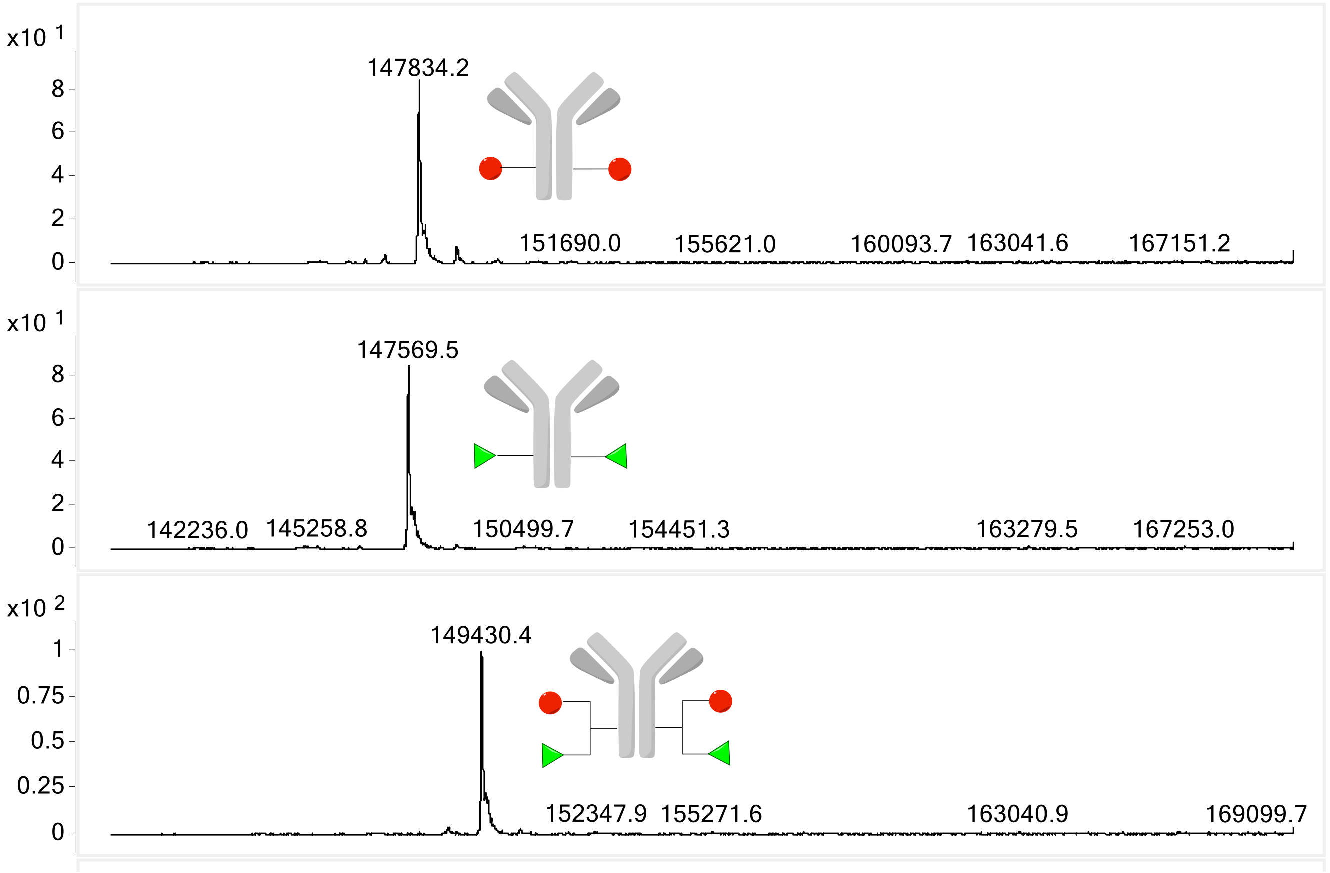

Deconvoluted ESI-mass spectra of N297A Trastuzumab–Cy5.5 conjugate (top), N297A Trastuzumab–AF48 conjugate (middle), and N297A Trastuzumab–AF488/Cy5.5 dual conjugate (bottom).

#### 3. REFERENCES

1. Anami *et al.* Glutamic acid–valine–citrulline cleavable linkers ensure high stability and efficacy of antibody–drug conjugates in mouse models. *Nat. Commun.*, **9**:2512 (2018).
2. Shi, Y. *et al.* Engagement of immune effector cells by trastuzumab induces HER2/ERBB2 downregulation in cancer cells through STAT1 activation. *Breast Cancer Res.* **16,** R33 (2014).
3. Anami, Y. *et al.* Enzymatic conjugation using branched linkers for constructing homogeneous antibody-drug conjugates with high potency. *Org. Biomol. Chem.* **15,** 5635–5642 (2017).
